# An ancestral tRNA maturation enzyme repurposed for cytoplasmic non-coding RNA processing

**DOI:** 10.64898/2026.10.07.757187

**Authors:** Athanasios-Nasir Shaukat, Mahmoud F. Dondeti, Alexandros Maniatis, Ilias Skeparnias, Eleni G. Kaliatsi, Dimitrios G. Anastasakis, Jinwei Zhang, Anastasios Vourekas, Constantinos Stathopoulos

**Author notes:** Correspondence to: Constantinos Stathopoulos.

## Abstract

Ribonuclease Z enzymes remove 3′ trailers from precursor tRNAs. In human cells, ELAC2/RNase Z^L^ matures nuclear and mitochondrial pre-tRNAs, whereas the cytosolic short paralog ELAC1/RNase Z^S^ repairs ANKZF1-cleaved tRNAs, but its broader substrate repertoire remains unclear. We investigated ELAC1 function using transcriptome profiling, small RNA sequencing, CLIP-seq, and biochemical reconstitution. ELAC1 was dispensable for canonical tRNA processing but associated with structured RNA polymerase III transcripts, prominently Y RNAs and vault RNAs (vtRNAs). Purified ELAC1 cleaved all four human vtRNAs near their 3′ ends, with high-resolution mapping identifying a conserved cleavage position within the 3′ extension. Efficient cleavage required structural features of the vtRNA terminal region. In contrast, ELAC2 showed little or no vtRNA cleavage under the conditions tested and displayed stricter substrate and cleavage-site selection on pre-tRNA. Loss and re-expression of ELAC1 altered structured ncRNA abundance and small-RNA end profiles. These findings identify vtRNAs as direct ELAC1 substrates and support functional specialization of human ELAC paralogs, with ELAC2 favoring precise pre-tRNA maturation and ELAC1 exhibiting broader RNA substrate specificity.

## Introduction

Ribonuclease Z (RNase Z) is a conserved endoribonuclease that catalyzes the removal of 3′ trailer sequences from precursor tRNAs, a fundamental step in tRNA maturation across all domains of life (1). In eukaryotes, this activity is carried out by two paralogs: a long form, RNase Z^L^ (ELAC2 in humans), and a short form, RNase Z^S^ (ELAC1 in humans) (Figure 1A) (2). ELAC2 is essential for cell viability and serves as the primary enzyme for nuclear and mitochondrial pre-tRNA processing, exhibiting high catalytic efficiency and strict substrate specificity (3–6). ELAC1, by contrast, represents the ancestral variant, is predominantly localized to the cytoplasm, and is dispensable for cell viability suggesting that it has diverged from canonical tRNA biogenesis and may fulfill alternative cellular roles (5).

**Figure 1.**
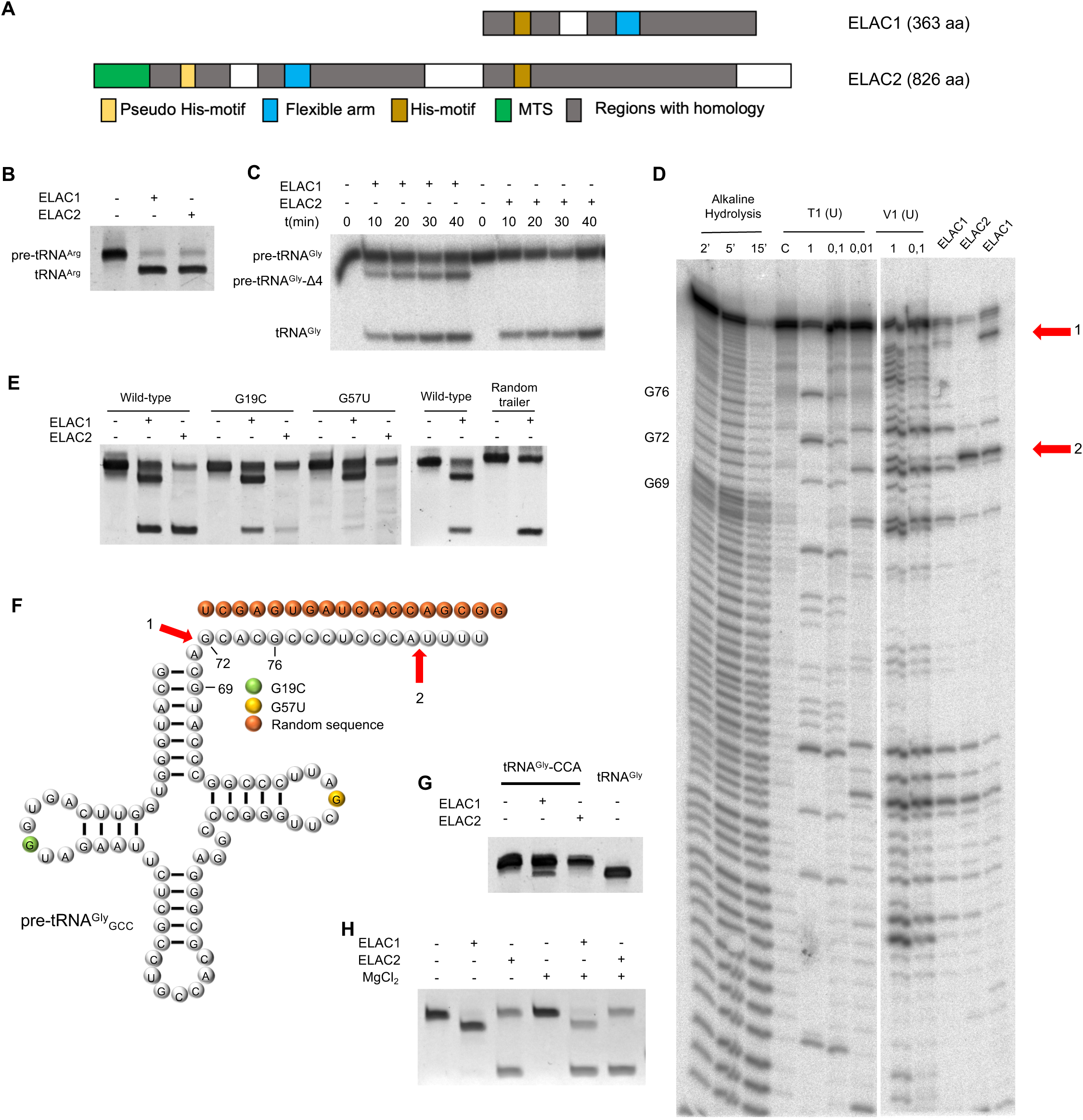
Human RNase Z paralogs differ in domain organization and in their structural requirements for pre-tRNA processing. (A) Domain organization of human ELAC1 (363 aa) and ELAC2 (826 aa), aligned on the catalytic His motif; regions of homology between the two proteins are shown in grey. ELAC2 additionally carries a mitochondrial targeting sequence and, in its N-terminal pseudo-nuclease half, a pseudo-His motif and the flexible arm, whereas in ELAC1 the arm lies C-terminal to the His motif. (B) In vitro cleavage of pre-tRNA^Arg^ (*D. discoideum*) by recombinant ELAC1 and ELAC2. (C) Time-course analysis of cleavage of 5′-^32^P-labeled pre-tRNA^Gly^_GCC_ by ELAC1 and ELAC2. (D) Structural probing of pre-tRNAGlyGCC by RNase T1, RNase V1 and alkaline hydrolysis; guanosines are numbered on the RNase T1 lanes. G76, within the 3′ trailer, lies in a V1-reactive double-stranded region. (E) Cleavage assays of pre-tRNA^Gly^_GCC_ wild-type, G19C, G57U and random-trailer variants by ELAC1 and ELAC2. (F) Schematic of the pre-tRNA^Gly^_GCC_ variants and ELAC1 cleavage sites. (G) Cleavage of mature tRNA^Gly^-CCA by ELAC1 and ELAC2. (H) Cleavage of pre-tRNA^Gly^_GCC_ by ELAC1 and ELAC2 in the presence or absence of Mg^2+^.

Biochemical and structural studies support this functional divergence. Although both enzymes recognize tRNA substrates, ELAC1 displays markedly reduced catalytic efficiency toward canonical pre-tRNAs compared to ELAC2, indicating that it is not optimized for this role (3, 7). Instead, ELAC1 appears to specialize in the removal of 2′-3′ cyclic phosphates from tRNAs generated during ribosome-associated quality control pathways (7). However, this activity in human cells is not unique to ELAC1, as ANGEL2 has recently been shown to perform the same repair function, suggesting that cyclic phosphate removal may not be a defining feature of ELAC1 biology (8). These observations hint at a role for ELAC1 in cytoplasmic RNA surveillance beyond tRNA repair.

Transcriptome-wide sequencing has revealed that a large fraction of the human genome is transcribed into non-coding RNAs (ncRNAs), many of which adopt defined secondary structures and function within ribonucleoprotein complexes (9). A substantial proportion of these ncRNAs are produced by RNA polymerase III, including tRNAs, Y RNAs, vault RNAs (vtRNAs), small nuclear RNAs (snRNAs), and small nucleolar RNAs (snoRNAs) (9, 10). These transcripts are not functionally uniform. tRNAs, snRNAs, and snoRNAs have long-established roles in translation and RNA processing, whereas the cellular functions of Y RNAs and vtRNAs are less firmly established. Members of both groups have been linked to the control of gene expression, cellular stress responses, and innate immunity (11–13). Notably, many RNA polymerase III transcripts share common structural features like stable stem-loop architectures and 3′ oligo(U) extensions, that influence RNA stability, localization, and protein interactions (9). It has also become increasingly clear that structured ncRNAs give rise to diverse populations of small RNA fragments with regulatory functions. Beyond well-characterized classes such as microRNAs (miRNAs), small interfering RNAs (siRNAs), and PIWI-interacting RNAs (piRNAs), additional small RNAs originate from tRNAs, Y RNAs, vtRNAs, rRNAs, snRNAs, and snoRNAs. These include tRNA-derived fragments (tRFs), Y RNA-derived RNAs, and vtRNA-derived small RNAs, which have been implicated in processes ranging from translational control to stress responses and antiviral defense (14, 15). One of these classes, the tRF-1 species, is released from the 3′ trailer during pre-tRNA maturation and has been attributed to RNase Z activity (16), which makes the trailer-processing enzymes candidates for their production. Despite their emerging importance, the enzymatic pathways responsible for generating and regulating these RNA fragments remain incompletely understood, particularly for RNA polymerase III-derived transcripts.

Two of these classes are particularly relevant here. vtRNAs and Y RNAs both fold into defined terminal stems that end in single-stranded 3′ extensions, an architecture that resembles the acceptor stem and 3′ trailer of a precursor tRNA. vtRNAs have been implicated in apoptosis, autophagy, and viral infection, partly through their capacity to generate regulatory RNA fragments or to modulate RNA-binding proteins (17). Y RNAs participate in ribonucleoprotein assembly and have been identified as endogenous ligands of innate immune sensors such as RIG-I (18). In both cases the reported activities depend on the structure and processing state of the RNA, which places their 3′ ends in a position to influence function.

Enzymes that evolved to act on pre-tRNAs could in principle recognize these features. Several ncRNAs harbor tRNA-like elements, including acceptor stem-like helices and accessible 3′ termini, that could serve as a basis for recognition by RNase Z enzymes (19, 20). Of the two human paralogs, ELAC1 is the more plausible candidate: its relaxed specificity for canonical pre-tRNAs and its cytoplasmic localization place it where mature vtRNAs and Y RNAs accumulate. This predicts that ELAC1, rather than ELAC2, acts on the 3′ ends of structured RNA polymerase III transcripts.

This prediction has not been tested. The physiological substrates of ELAC1 and its contribution to ncRNA metabolism remain poorly characterized, and it is unclear whether ELAC1 directly processes RNA polymerase III transcripts, contributes to small RNA biogenesis, or functions in cytoplasmic RNA quality control pathways that influence RNA stability and ribonucleoprotein assembly. To address these questions, we investigated the role of ELAC1 in human cells using an integrated approach combining transcriptome profiling, small RNA sequencing, CLIP-seq, and *in vitro* biochemical assays. ELAC1-associated RNA profiles revealed strong enrichment for RNA polymerase III transcripts, including vtRNAs and other structured ncRNAs. Disruption and re-expression of ELAC1 were accompanied by widespread changes in ncRNA abundance and in derived small RNA populations. Purified ELAC1 directly cleaved all four tested human vtRNA transcripts, whereas ELAC2 showed little activity on these substrates under the conditions tested. Together, these findings identify vtRNAs as direct biochemical ELAC1 substrates and provide a testable model of RNase Z substrate partitioning.

## Materials and Methods

### Cell lines and culture conditions

HEK-293T cells (ATCC CRL-3216) were kindly provided by Dr. Dionysios Chartoumpekis (University of Patras) and maintained in Dulbecco’s Modified Eagle Medium (DMEM) supplemented with 10% fetal bovine serum (FBS) at 37°C in a humidified incubator with 5% CO₂, and were used between passages 4 and 12. Cultures tested negative for mycoplasma. The ELAC1 knockout cell line was generated by CRISPR-Cas9-mediated targeting of the second exon of the *ELAC1* gene using the pSpCas9(BB)-2A-Puro (PX459) V2.0 vector (Addgene plasmid #62988; RRID:Addgene_62988) (21). sgRNA and genotyping primer sequences are listed in Supplementary Table S1. Following puromycin selection, clonal isolation was performed by serial dilution. Successful disruption of *ELAC1* was confirmed by Sanger sequencing, which identified a frameshift-inducing 22 bp deletion in the targeted exon. Attempts to generate an *ELAC2* knockout using the same approach were unsuccessful (data not shown), consistent with its essential role in tRNA maturation. For rescue and overexpression experiments, cells were transiently transfected with pcDNA3.1-ELAC1 or empty pcDNA3.1 vector using polyethylenimine (PEI) and harvested 48 hours post-transfection.

### Western blotting

Whole-cell lysates from wild-type and ELAC1 knockout HEK-293T cells were resolved by SDS-PAGE on 12% polyacrylamide gels using 2× Laemmli buffer (25% v/v glycerol, 5% β-mercaptoethanol, 2% w/v SDS, 0.01% w/v bromophenol blue, 65 mM Tris pH 6.8) and heating to 95°C for 10 minutes, in Tris/glycine/SDS running buffer at 120 V. Proteins were transferred to a PVDF membrane (Millipore) at 42 V for 3 hours at 4°C, and membranes were blocked for 3 hours at room temperature in 5% non-fat milk in TBS. Membranes were incubated with anti-ELAC1 primary antibody (1:100 in 3% non-fat milk-TBS) for 3 hours at room temperature or overnight at 4°C, washed three times in TBS, and incubated with HRP-conjugated anti-mouse secondary antibody (1:1000) for 1 hour at room temperature. Bands were visualized using an ECL detection kit (Bio-Rad) and X-ray film (Santa Cruz Biotechnology) (antibody details in Supplementary Table S7).

### Cell cycle analysis

HEK-293T and knockout cells were seeded in 6-well plates and synchronized by serum starvation for 24h. Cells were subsequently released into the cell cycle by the addition of complete medium and harvested for 24 h. Then, the cells were fixed in ice-cold 70% ethanol overnight, washed with PBS, and stained with propidium iodide (10 μg/ml) and RNase A (100 μg/ml) at 37°C for 30 min. DNA content was analyzed using a BD Biosciences BD Accuri C6 Plus Flow Cytometer. Data analysis was performed in FlowJo v10.0.7.

### Recombinant protein expression and purification

The ELAC1 and ELAC2 coding sequences were obtained from GenScript in pcDNA3.1 vectors, and the ELAC1 catalytically inactive H64A mutant used for CLIP-seq and biochemical assays was generated from this construct by site-directed mutagenesis (Supplementary Table S3). Full-length ELAC1 and ELAC2 coding sequences, including the stop codon, were subcloned from these pcDNA3.1 constructs into pET-28b(+) using NdeI and XhoI restriction sites (primers listed in Supplementary Table S2). Recombinant His-tagged ELAC1 and ELAC2 were expressed in *E. coli* BL21(DE3) Rosetta2 pLysS cells from pET-28b(+) constructs. Cultures were grown at 37°C to OD₆₀₀ 0.4–0.5, induced with 0.5 mM IPTG, and incubated at 18°C for 16 hours. Cells were lysed by sonication in 50 mM Tris-HCl pH 7.5, 100 mM KCl, 5% glycerol, 10 mM imidazole. After clarification (10,000 × g, 4°C), the lysate was purified using 1 ml bed volume of Ni-NTA affinity chromatography resin (Macherey-Nagel) and eluted using 50 mM Tris-HCl pH 7.5, 100 mM KCl, 5% glycerol, 250 mM imidazole. Protein concentration was determined spectrophotometrically. Final polishing was performed by size exclusion chromatography on a Superdex 200 10/300 GL column (GE Healthcare). Purified proteins were stored in 20 mM Tris-HCl pH 7.5, 50 mM NaCl, 1.5 mM MgCl₂, 1 mM DTT, 20% glycerol at −80°C as single-use aliquots that were thawed once.

### *In vitro* transcription and 5′ end labeling

For each ncRNA gene, the proximal genomic locus was amplified and cloned into the pJET1.2/blunt vector using the CloneJET PCR Cloning Kit (Thermo Scientific); the resulting plasmids served as templates for subsequent PCR amplification (primers listed in Supplementary Table S2), with the exception of the *Dictyostelium discoideum* pre-tRNA^Arg^ template, which was synthesized and supplied in a pUC57 vector by GenScript. DNA templates for in vitro transcription were generated by PCR using Q5 High-Fidelity DNA Polymerase (NEB) from these plasmid constructs using the primer pairs listed in Supplementary Table S4. Each forward primer incorporates a T7 RNA polymerase promoter sequence. In vitro transcription reactions were carried out with T7 RNA polymerase (NEB) at 37°C for 16 hours in the presence of 2 mM each NTP. The DNA template was subsequently removed by DNase I (NEB) treatment, and RNA was purified by phenol/chloroform extraction followed by denaturing urea-PAGE. Transcripts were visualized by UV shadowing, excised, and eluted overnight at room temperature in 20 mM Tris-HCl pH 7.5, 300 mM sodium acetate, 1 mM EDTA, 0.25% SDS. Point substitutions and internal deletions in the vtRNA1-2, pre-tRNA^Gly^_GCC_ and pre-tRNA^Arg^ templates (including the G19C, G57U, random-trailer and acceptor-stem-length variants) were introduced into the corresponding pJET1.2 plasmids by site-directed mutagenesis (Supplementary Table S3); where a change fell within the first or last few nucleotides of the transcript, it was instead incorporated directly into the PCR primer used to generate that construct’s in vitro transcription template (Supplementary Table S4). All primer and construct sequences are provided in Supplementary Tables S1–S4. Eluted RNA was recovered by phenol/chloroform extraction and ethanol precipitation; the sequence of each in vitro transcribed RNA is provided in Supplementary Table S5.

For 5′ end labeling, transcripts were first dephosphorylated using Antarctic Phosphatase (NEB) at 37°C for 1 hour, followed by phenol/chloroform extraction and ethanol precipitation. The 5′ ends were then labeled using [γ-³²P]-ATP and T4 Polynucleotide Kinase (NEB, M0201S) at 37°C for 30 minutes. Radiolabeled transcripts were purified by denaturing urea-PAGE followed by autoradiography and gel excision.

For vtRNA1-2 deletion mutants generated by oligonucleotide hybridization (vtRNA mutant, VM, constructs), complementary oligonucleotides were dissolved in annealing buffer (10 mM Tris-HCl pH 8.0, 50 mM NaCl, 1 mM EDTA), hybridized by heating to 95°C followed by slow cooling to 25°C, and used directly as templates for in vitro transcription at a final concentration of 100 nM.

### *In vitro* cleavage assays

Standard cleavage assays were performed in 20 mM Tris-HCl pH 7.5, 50 mM NaCl, 1.5 mM MgCl₂, 1 mM DTT at 37°C for 10 minutes, with ELAC1 at 100 nM and ELAC2 at 10 nM unless otherwise stated. RNA substrates were refolded prior to addition of enzyme by heating to 65°C for 2 minutes followed by equilibration at 37°C for 5 minutes. Reactions were terminated by phenol/chloroform extraction and analyzed by denaturing urea-PAGE, stained with SYBR Gold and imaged on a ChemiDoc system (Bio-Rad). For assays using 5′-radiolabeled substrates, products were visualized by phosphor imaging on an FLA-3000 instrument (Fujifilm). Equal RNA mass was loaded in each lane and reaction volumes were equalized in 1× reaction buffer. For kinetic analysis of pre-tRNA^Arg^ cleavage, substrate concentrations of 25–500 nM were used at enzyme concentrations of 15 nM (ELAC1) and 0.1 nM (ELAC2), with reaction times of 2, 5, and 10 minutes. Initial velocities were fitted to the Michaelis-Menten equation using GraphPad Prism v8. For Mg²⁺-free conditions, MgCl₂ was omitted from the reaction buffer. Deviations from standard conditions are noted in the relevant figure legends.

### Enzymatic RNA sequencing and footprinting and Northern blotting

Enzymatic RNA sequencing and footprinting of 5′-radiolabeled RNA substrates was performed using RNase T1, RNase A, and RNase V1 under native and denaturing conditions as appropriate, following established protocols. Products were resolved by high-resolution denaturing urea-PAGE and visualized by phosphor imaging.

Total RNA (10 μg) was resolved on a 15% denaturing urea-PAGE gel in 0.5× TBE, stained briefly with ethidium bromide to verify equal loading, and transferred to a Hybond N+ nylon membrane (Amersham) at 250 mA for 1 hour. The membrane was UV-crosslinked at 254 nm for 60 seconds. DNA probes (10 pmol) were 5′-radiolabeled with [γ-³²P]-ATP using T4 PNK (NEB) at 37°C for 30 minutes and diluted directly into Church hybridization buffer (0.5 M sodium phosphate pH 7.2, 10 mg/ml BSA, 1 mM EDTA, 7% SDS). The membrane was pre-hybridized at 37°C for 3 hours, then hybridized with the radiolabeled probe at 37°C for 3 hours. Post-hybridization washes were performed sequentially with 2× SSC/0.05% SDS (twice, 30 minutes each) and 0.1× SSC/0.1% SDS (twice, 15 minutes each). The membrane was exposed to a Fujifilm phosphor imaging plate and scanned on an FLA-3000 phosphor imager. The probe sequences are listed in Supplementary Table S6).

### CLIP-seq

CLIP-seq was performed in HEK-293T cells expressing FLAG-tagged wild-type ELAC1 or the catalytically inactive H64A mutant, following an established protocol (22, 23). Cells were UV-crosslinked at 400 mJ/cm² on ice and lysed in buffer containing 2% Empigen with protease and RNase inhibitors and DNase. Crosslinked RNA-protein complexes were immunoprecipitated using M2 anti-FLAG magnetic beads (Sigma). cDNA libraries were sequenced on an Illumina HiSeq2500 platform. Reads were trimmed using Cutadapt v4.1 (24) and mapped to hg19 using STAR v2.5.4b (25). Genome coverage was normalized by counts per million (CPM) using bam Coverage from deep Tools v3.5.1 (26) and visualized in the UCSC Genome Browser.

### Whole transcriptome sequencing and analysis

Total RNA was isolated from HEK-293T cells using TRIzol reagent (Invitrogen) according to the manufacturer’s instructions. RNA was treated with DNase I (NEB) to remove genomic DNA contamination, followed by phenol/chloroform extraction and ethanol precipitation. RNA quantity and quality were assessed by spectrophotometry and agarose gel electrophoresis. For small RNA enrichment, the mirVana miRNA Isolation Kit (Invitrogen) was used to separate the >200 nt and <200 nt fractions. For the rescue experiment small RNA libraries, enrichment was performed by denaturing 15% urea-PAGE, with the 15–150 nt region excised, eluted overnight in gel elution buffer, and recovered by phenol/chloroform extraction and ethanol precipitation.

The >200 nt RNA fraction was ribodepleted and used for library preparation. For the knockout experiment, ribodepletion was performed using the NEBNext rRNA Depletion Kit (NEB) and libraries were prepared using the TruSeq RNA Library Prep Kit v2 (Illumina), followed by paired-end sequencing on a NovaSeq 6000 (2×50 cycles). For the rescue experiment, ribodepletion used the RiboMinus Eukaryote Kit v2 (Invitrogen) and libraries were prepared using the Ion Total RNA-seq Kit v2 (Ion Torrent), followed by single-end sequencing on an Ion GeneStudio S5 instrument. Reads from both platforms were trimmed with fastp v0.23.2 (27) or Trimmomatic v0.38 respectively (28), mapped to hg38 using STAR v2.7.8a (25), and quantified using featureCounts v2.0.1 with Ensembl annotation (GCA_000001405.28) (29). Differential expression analysis was performed using limma-voom v3.50.1, with a minimum expression threshold of CPM >0.5 in at least one sample (30). Gene Ontology Biological Process and Reactome overrepresentation analysis of the differentially expressed genes used the PANTHER Overrepresentation Test (31) (release 20221013) with Fisher’s exact test and Benjamini–Hochberg correction, against the *Homo sapiens* reference list of 20,589 genes (GO release 2023-03-06; Reactome version 77 (32)); 496 of the submitted genes mapped to PANTHER entries.

### Small RNA sequencing and analysis

Small RNAs were enriched from total RNA and size-fractionated into 15–50 nt and 50–100 nt fractions as described above. For the knockout and overexpression libraries, small RNA libraries were prepared using a ligation-based protocol with Rnl2(1–249) K227Q ligase for 3′ adapter ligation and T4 Rnl1 for 5′ adapter ligation, followed by reverse transcription using SuperScript IV (Thermo Fisher) and PCR amplification. Size selection was performed using a 3% agarose Pippin Prep system and libraries were sequenced on an Illumina HiSeq4000. For the rescue experiment, libraries were prepared using the Ion Total RNA-seq Kit v2 (Ion Torrent) with modifications to preserve both tRNA and smaller RNA species (adapter hybridization at 70°C; ligation at 16°C for 16 hours; reverse transcription at 57°C for 90 minutes; size-selection step omitted) and sequenced on an Ion GeneStudio S5 instrument. Because the knockout and overexpression libraries and the rescue libraries were therefore produced on different platforms and library workflows, the three comparisons were analyzed as independent comparisons rather than as a single matched series. Reads were separated into 15–50 nt and >50 nt fractions using Trimmomatic v0.38 (28). The >50 nt fraction was mapped to hg38 using Bowtie2 v2.4.5 (33) (--very-sensitive-local) and quantified using featureCounts v2.0.1 with Ensembl annotation (34). For tRNA analysis, reads were mapped to a custom artificial chromosome containing all nuclear and mitochondrial tRNA sequences from gtRNAdb (35), tRNA-derived fragments were annotated using MINTmap v2.0 (36) and named according to the tDRnamer nomenclature (37). miRNAs were annotated using miRBase v22 (38). Additional small RNA classes, including snRNA-derived RNAs (snDRs), snoRNA-derived RNAs (snoDRs), rRNA-derived RNAs (rDRs), Y RNA-derived RNAs (yDRs), and vtRNA-derived RNAs (vtDRs), were annotated using SPORTS1.1 (39). Differential expression analysis across all small RNA classes was performed using limma-voom v3.50.1 (30) with CPM >0.5 in at least one sample as the minimum expression threshold.

## Results

### ELAC1 and ELAC2 recognize related RNA substrates but differ in structure-dependent cleavage-site selection

ELAC1 and ELAC2 share the RNase Z catalytic core but differ in domain organization (Figure 1A). To establish how this divergence relates to substrate recognition, and how the two enzymes differ mechanistically, we first examined the processing of pre-tRNA substrates with distinct structural properties. Both enzymes cleave the pre-tRNA^Arg^ (from *Dictyostelium discoideum*) at the canonical site immediately downstream of the discriminator base (Figure 1B, Supplementary Figure S1D). Both therefore recognize canonical pre-tRNA geometry. ELAC2 is markedly more efficient than ELAC1 on this substrate (3). In our measurements the difference in apparent catalytic efficiency is close to two orders of magnitude (Figure S1A,B,C). The assay uses three time points and does not saturate ELAC1, so this is best read as an order-of-magnitude difference rather than a precise ratio. Substrate recognition is conserved, but ELAC2 is optimized for rapid processing of pre-tRNAs destined for translation.

The two enzymes diverge substantially on pre-tRNA^Gly^_GCC_. ELAC2 cleaves exclusively at the canonical site, whereas ELAC1 generates an additional cleavage within the 3′ trailer (Figure 1C, F). Enzymatic RNA sequencing and footprinting localized this secondary site just upstream of the poly(U) tract (Figure 1D). RNase V1 cleavage in this region indicates that the 3′ trailer is base paired rather than single stranded. Together with the native PAGE behavior described below, we take this as evidence that the trailer can adopt an alternative pairing, whose structure we have not determined. Across the rest of the molecule the V1 pattern matches the known secondary structure of the substrate, with additional paired positions attributable to the coexistence of the two conformations, and G76, the last guanosine within the trailer, falls clearly inside a V1-reactive double-stranded region. Native PAGE showed that pre-tRNA^Gly^_GCC_ adopts two conformations in the presence of Mg²⁺ and a single resolved species without added Mg²⁺ (Supplementary Figure S1F). Under the same conditions ELAC1 shifts its cleavage preference toward the upstream trailer site, while ELAC2 continues to generate mature tRNA (Figure 1H). Both enzymes carry a buried catalytic zinc acquired during folding, so omitting Mg²⁺ alters the fold of the substrate without affecting catalysis.

The simplest account of these data is that the two conformations do not interconvert appreciably over the course of the reaction. ELAC2 acts only on the canonically folded molecules and therefore leaves the alternative conformer uncleaved, which accounts for the larger residual substrate in the ELAC2 lanes, whereas ELAC1 cleaves the alternative conformer and consumes more of the population. Consistent with this, the intensity of the alternative cleavage product relative to mature tRNA approximately matches the ratio of the two species on the native gel. Were the two conformations in rapid equilibrium, ELAC2 would be expected to deplete the substrate regardless of the starting distribution, which is not what we observe.

Canonical cleavage by both enzymes depends on the tertiary elbow. G57U abolished production of mature tRNA by ELAC1 and ELAC2, and G19C reduced it to a residual level, whereas the ELAC1-specific trailer cleavage proceeded efficiently in both mutants (Figure 1E,F). The two cleavage events therefore read different structural features: the canonical reaction requires the folded elbow, while the secondary reaction requires only the trailer duplex. The magnesium dependence separates the two enzymes rather than the two sites. ELAC2 generates mature tRNA without added Mg²⁺ at a yield comparable to the standard reaction, so its canonical activity does not require magnesium-dependent compaction of the substrate. ELAC1, whose canonical activity is weak to begin with, produces only the trailer product under the same conditions. The simplest reading is that ELAC1 requires the magnesium-stabilized state to cleave canonically at all, while its activity at the trailer duplex is unaffected, leaving it with a single detectable product.

The dependence of the secondary cleavage on the 3′ trailer sequence of pre-tRNA^Gly^_GCC_ was confirmed by replacing it with a randomly generated sequence, which abolished the additional cleavage while leaving canonical processing intact (Figure 1E). This indicates that the native trailer sequence is required for the secondary cleavage. That cleavage has so far been observed only in vitro. In the cellular small RNA data, pre-tRNA fragments ending within the 3′ trailer terminate overwhelmingly inside the genomically encoded U-run rather than upstream of it, and the minority ending upstream are not enriched in ELAC1 knockout cells, so we find no evidence that the secondary cleavage occurs in cells. Altering the acceptor stem length reduced canonical processing by both enzymes but did not abolish the ELAC1-specific secondary cleavage (Figure S1E). Acceptor stem length is therefore a recognition determinant for canonical processing but not for the non-canonical activity. Finally, ELAC1 retained the ability to cleave tRNA^Gly^-CCA, albeit at reduced efficiency, while ELAC2 was inhibited by the CCA sequence, consistent with its known role in preventing aberrant cleavage of mature tRNAs (Figure 1G). Collectively, these results indicate that ELAC2 has evolved high fidelity and efficiency on pre-tRNA substrates. ELAC1 instead acts on structurally heterogeneous or non-canonically folded RNAs, which is consistent with its broader substrate repertoire. One possible explanation is that ELAC1 has shifted from reading the tRNA fold to reading a duplex-to-single-strand junction, perhaps with a preference for a 3′ U-rich tail. Our data do not distinguish this from the alternatives.

### CLIP-seq reveals broad association of ELAC1 with RNA polymerase III transcripts

To identify RNA substrates of ELAC1 *in vivo*, we performed CLIP-seq in HEK293T cells expressing either wild-type ELAC1 or a catalytically inactive mutant (H64A). The mutant was intended to trap substrates before cleavage and to capture transient interactions missed by the wild-type enzyme. ELAC1 associates with a diverse set of transcripts strongly enriched for RNA polymerase III products, including tRNAs, snoRNAs, snRNAs, Y RNAs, vtRNAs, and specific lncRNAs such as RPPH1, the catalytic RNA subunit of RNase P (Figure 2A, S2A). tRNAs recovered in the CLIP data were predominantly mature, with limited coverage in the 3′ trailer regions, consistent with a role in cytoplasmic tRNA surveillance and repair, and not in primary pre-tRNA processing. Among snRNAs, U6, the only RNA polymerase III-transcribed spliceosomal snRNA, was identified alongside a subset of non-intronic snoRNAs, pointing to a potential role for ELAC1 in their cytoplasmic processing or quality control. Reads mapping to RPPH1, RMRP, and 7SK were concentrated over short regions rather than distributed across the full-length transcripts (Figure S2A). Because unbound RNA is degraded by the RNase treatment during the CLIP procedure, this localized coverage is best read as a binding footprint rather than as evidence that ELAC1 binds fragments generated in cells.

**Figure 2.**
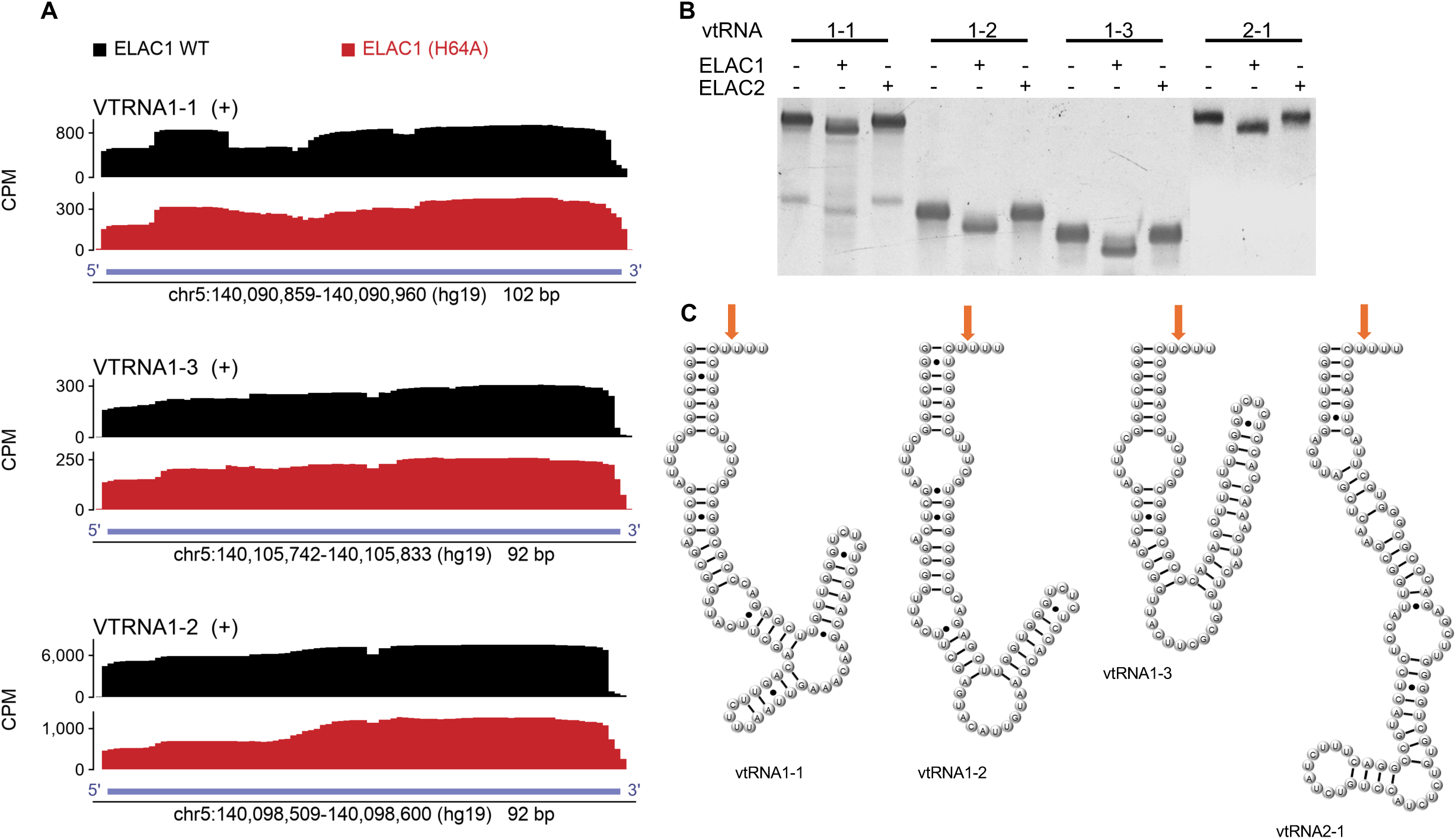
ELAC1 associates with vault RNA loci in cells and cleaves their 3′ ends in vitro. (A) CLIP-seq read coverage of ELAC1-WT and the catalytic mutant ELAC1-H64A across the human vtRNA1 loci: per-base sense-strand coverage in counts per million, with the window coordinates (hg19) and the gene body drawn to scale beneath each track pair. Mean coverage across the plotted window is 740 versus 293 CPM at VTRNA1-1 (WT/H64A 2.5-fold), 6,252 versus 933 CPM at VTRNA1-2 (6.7-fold) and 245 versus 213 CPM at VTRNA1-3 (1.2-fold); VTRNA2-1 is not shown because coverage there is too low to measure (0.2 versus 0.06 CPM). One library per genotype. (B) In vitro cleavage assays of the four human vault RNAs using recombinant ELAC1 (100 nM) or ELAC2 (10 nM) at 37°C for 10 min. Unlabeled transcripts, visualized by SYBR Gold staining. (C) Predicted secondary structures of the four human vault RNAs.

vtRNAs were the most abundantly represented non-tRNA class in the CLIP data. All three vtRNA1 family members showed increased read coverage at their 3′ ends in the catalytic mutant relative to wild-type, a pattern expected if the mutant traps substrates at the site of cleavage (Figure 2A). Their predicted tRNA-like acceptor stem fold provides a potential structural basis for ELAC1 recognition. Three observations collectively support this conclusion: vtRNA enrichment, structural similarity to the tRNA acceptor stem, and the differential 3′-end coverage between wild-type and mutant. Together these make vtRNAs the strongest non-tRNA substrate candidates for ELAC1 in the cytoplasm, which we test directly by biochemical reconstitution below. All four Y RNAs were also recovered in the CLIP data and, like vtRNAs, adopt stem-loop structures resembling the tRNA acceptor stem. A modest increase in coverage at the 3′ single-stranded U-rich tract of RNY4 was observed in the catalytically inactive mutant relative to wild-type, hinting that this region may represent a cleavage site (Figure S2A).

### ELAC1 specifically cleaves the 3′ extension of vault RNAs

To biochemically validate vtRNAs as direct ELAC1 substrates and define the cleavage site, all four vtRNA transcripts were tested in cleavage assays with recombinant ELAC1 and ELAC2 (Figure 2B). ELAC1 cleaved all four vtRNAs, while ELAC2 showed no detectable activity, an observation that indicates a strong substrate preference for ELAC1, but not absolute paralog exclusivity. Enzymatic RNA sequencing and footprinting of 5′-radiolabeled vtRNA1-2 placed the cleavage site immediately after U86, the first nucleotide of the four-nucleotide single-stranded tail that follows the terminal stem (Figure 3A). The other three paralogs are predicted to share this architecture, with the terminal stem closing on a cytidine in each, placing the equivalent residues at U96 in vtRNA1-1, U86 in vtRNA1-3 and U98 in VTRNA2-1. Cleavage at these positions releases a trinucleotide from each transcript, consistent with the products obtained from all four (Figure 2B, C), and indicates that the terminal-stem register rather than the local sequence sets the site.

**Figure 3.**
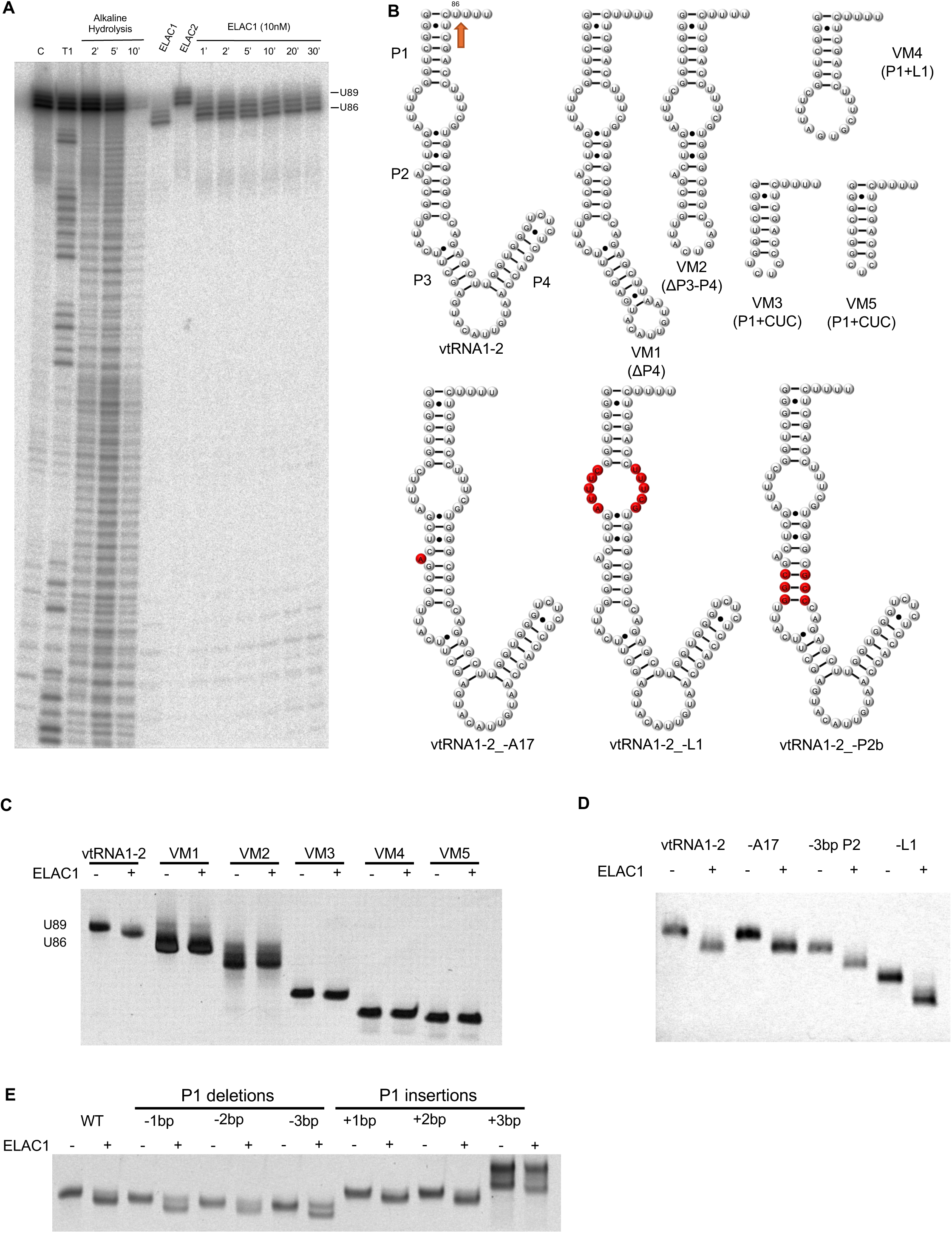
Structural determinants of ELAC1-mediated cleavage of vtRNA1-2. (A) Enzymatic footprinting of 5′-32P-labeled vtRNA1-2 identifying the ELAC1 cleavage site downstream of U86, visualized by phosphor imaging. (B) Secondary structure models of the vtRNA1-2 deletion and truncation variants, designated VM for vtRNA mutant. (C–E) In vitro cleavage assays of the variants drawn in (B): apical stem deletions (C), smaller internal deletions (D) and altered P1 stem lengths (E). Unlabeled transcripts, visualized by SYBR Gold staining. In the +3 bp lanes the upper band is a longer by-product of in vitro transcription that could not be removed; the lower band is the +3 bp transcript.

To identify structural elements required for ELAC1 recognition, we generated a series of deletion and truncation variants of vtRNA1-2 and tested them under identical assay conditions (Figure 3B). Sequential removal of the apical stems revealed that deletion of the P4 stem alone was sufficient to abolish cleavage, despite its distal position relative to the cleavage site (Figure 3C). Whether P4 contacts the enzyme directly or instead participates in tertiary interactions such as a pseudoknot that maintain the architecture required for ELAC1 engagement, remains to be determined. In contrast, deletion of the unpaired base A17, loop L1, and the P2b stem had no effect on cleavage efficiency or site specificity, demonstrating that these elements are dispensable for recognition (Figure 3D). Altering the P1 stem length produced an asymmetric effect. Shortening it by two or three base pairs reduced processing, seen as accumulation of unprocessed substrate, which could have been caused by end fraying and instability of the shorter stem. Lengthening was well tolerated. Neither modification appeared to alter the cleavage position, suggesting that variations in P1 stem length do not affect recognition specificity (Figure 3E). Together, these results show that ELAC1 cleaves vtRNA1-2 at a single position that none of the cleaved variants displaced, and that the elements required for cleavage are not confined to the region surrounding the site, since the distal P4 stem is essential while A17, L1, and P2b are dispensable.

### ELAC1 exhibits distinct cleavage patterns on non-tRNA substrates

Having established vtRNAs as direct ELAC1 substrates, we extended the *in vitro* cleavage assays to a broader panel of structured ncRNAs identified in the CLIP-seq data, including several RNA polymerase III transcripts. This tested the scope of ELAC1 activity and its specificity relative to ELAC2 (Figure S2B).

The results revealed three categories of substrate behavior. Two RNAs were cleaved by both enzymes but with distinct patterns. ELAC1 and ELAC2 both cleaved RMRP near its 5′ and 3′ ends, with ELAC1 generating an additional shorter fragment. Both also cleaved 7SK, with ELAC2 producing additional smaller products. RNAs cleaved exclusively by ELAC1 included RNU6 and RNY4, placing them in the same category as vtRNAs as ELAC1-specific substrates. SNORA73B gave a differential profile with ELAC1 cleaved near the transcript termini, producing a slight size shift, whereas ELAC2 cleaved internally to give shorter fragments. RNU2 was cleaved predominantly by ELAC1, yielding two principal smaller fragments. Finally, RPPH1 and SNORD78 were resistant to cleavage by either enzyme under the tested conditions. To confirm that ELAC1 targets the 3′ ends of these substrates, cleavage assays were performed using 5′-32P-labeled transcripts of RMRP, 7SK, RNU2, RNU6 and RNY4 (Figure S2C). Processing close to the 3′ end was observed for RMRP, 7SK, RNU6 and RNY4. Additional upstream sites were detected for RMRP, 7SK and RNU2, which may reflect recognition of internal stem-loop structures within these more complex transcripts by ELAC1. These findings show that ELAC1 processes a defined set of structured non-tRNA substrates with a product pattern distinct from ELAC2. Among the non-tRNAs tested, vtRNAs, RNY4 and RNU6 were cleaved by ELAC1 alone.

### Loss of ELAC1 does not alter tRF-1001 levels but broadly reprograms gene expression

Using CRISPR-Cas9, we targeted the second exon of the ELAC1 gene in HEK293T cells (Figure 4A). ELAC1 mRNA levels are essentially unchanged in the knockout (log2 fold change −0.16, adjusted P = 0.16, total RNA-seq), so the exon 2 disruption is a protein-level knockout that the RNA-seq does not independently confirm. Isolated knockout cells were viable but exhibited mild growth defects, including a reduced proliferation rate and an increased proportion of cells in the G1 phase (Figure 4A, S3A–C). ELAC2 is the canonical nuclear RNase Z responsible for pre-tRNA 3′ processing and tRF-1 biogenesis. However, earlier fractionation studies reported cytoplasmic accumulation of tRF-1 species, raising the possibility that the cytoplasmic paralog ELAC1 might contribute to their production. To test this directly, we analyzed tRF-1001, a well-characterized tRF-1 derived from tRNA^Ser(UGA)^, by northern blot in total RNA from wild-type, *ELAC1* knockout, and *ELAC1*-depleted HEK293T cells (Figure 4B). tRF-1001 levels were unchanged across all conditions, indicating that ELAC1 is dispensable for its generation.

**Figure 4.**
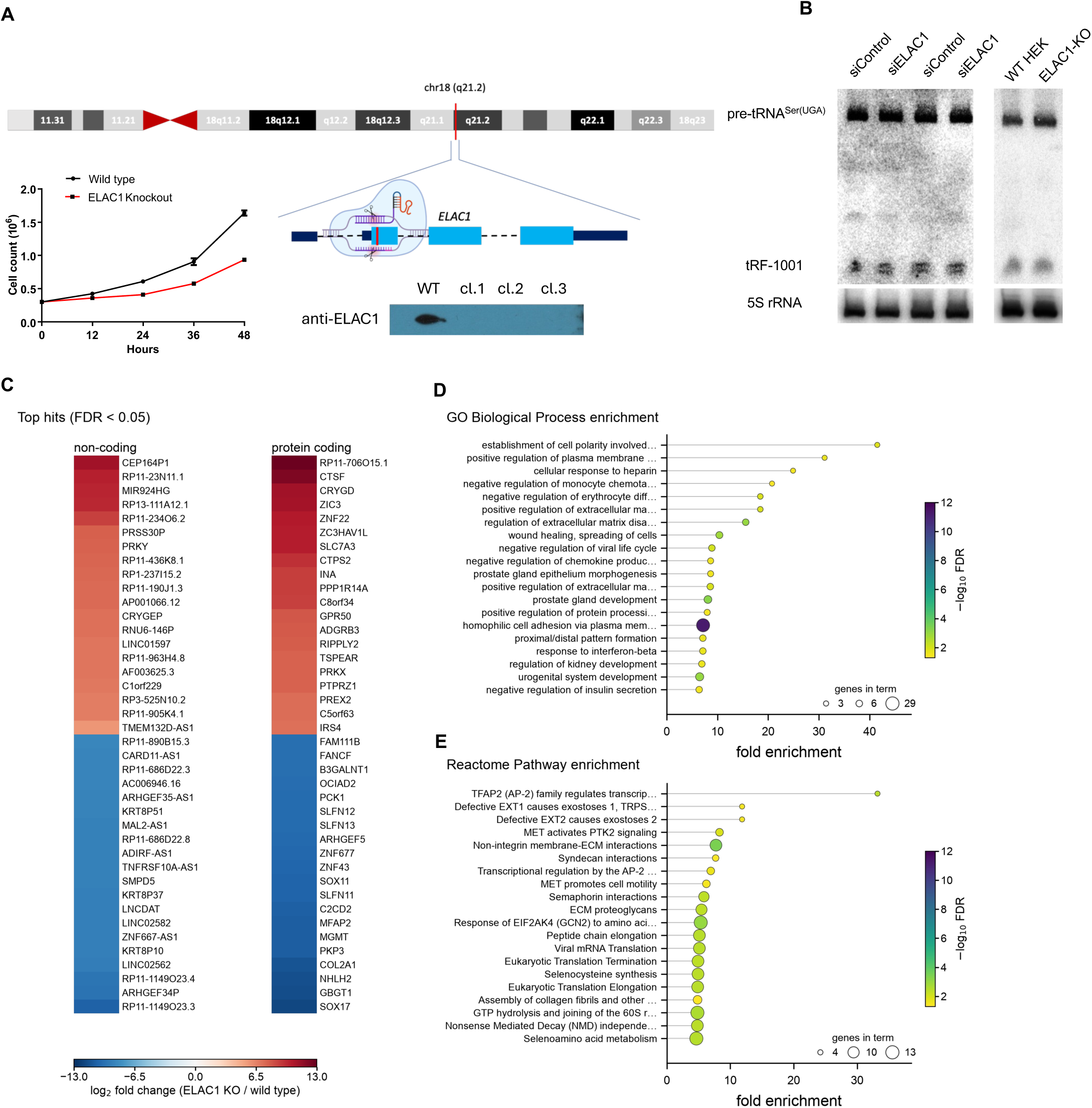
CRISPR–Cas9-mediated disruption of ELAC1 does not affect tRF-1001 biogenesis but broadly remodels the HEK293T transcriptome. (A) Schematic of the human ELAC1 locus (chromosome 18q21.2) and CRISPR–Cas9 knockout strategy. Immunoblot showing ELAC1 expression in parental cells and independent knockout clones (4, 5 and 7), and growth curves of wild-type and ELAC1 knockout cells over 48 h (n = 3). (B) Northern blot analysis of tRF-1001 and its precursor in siRNA-mediated knockdown and CRISPR knockout cells. 5S rRNA serves as the loading control. (C) Differentially expressed genes with the largest fold changes in ELAC1 knockout relative to wild-type cells (FDR < 0.05), shown as the 20 most strongly increased and 20 most strongly decreased loci within the non-coding (left) and protein-coding (right) sets; cell color gives log2 fold change on a diverging scale centered at zero. (D, E) Overrepresentation analysis of the differentially expressed gene list against GO Biological Process (D) and Reactome pathways (E): x-axis, fold enrichment; point area, number of input genes in the term; color, −log10 FDR. n = 2 biological replicates per condition; statistical procedures are given in Methods.

To characterize the broader impact of ELAC1 loss on the cellular transcriptome, we performed whole-transcriptome RNA-seq (>200 nt) on knockout cells, capturing mRNAs, long non-coding RNAs, and other RNA species. *ELAC1* knockout caused widespread changes in gene expression, suggesting broad transcriptome reprogramming (Figure S3D,E). Gene Ontology (GO) enrichment analysis of the 500 most significant differentially expressed genes by adjusted *P* value, of which 496 mapped to annotated PANTHER entries revealed significant effects on cell migration, cell adhesion, chemotaxis, extracellular matrix organization, and the negative regulation of viral replication and insulin secretion (Figure 4C–E).

### Rescue of ELAC1 expression in knockout cells leads to upregulation of non-coding RNAs

Throughout, rescue refers to transient expression of pcDNA3.1-ELAC1 in the ELAC1 knockout background and overexpression to the same construct in wild-type cells, each compared with the empty vector in the matching background and harvested 48 h after transfection. To examine how ELAC1 affects cellular RNA homeostasis, we reintroduced ELAC1 into knockout cells and compared the resulting transcriptome to that of knockout cells carrying an empty vector. Rather than restoring gene expression to a wild-type profile, ELAC1 re-expression caused a pronounced upregulation of multiple non-coding RNA classes, with miscRNAs representing the largest fraction, followed by lncRNAs, snRNAs, and snoRNAs (Figure 5A,B). This comparison is rescue against empty-vector knockout rather than against cells that underwent the same editing procedure with ELAC1 intact, and the transgene is expressed far above endogenous levels (below). It therefore cannot distinguish a genuine failure to rescue from effects of the knockout clone or of supraphysiological ELAC1. Several RNA polymerase III-derived transcripts are encompassed in the miscRNA group, including 7SK, 7SL and Y RNAs, suggesting that ELAC1 preferentially influences this class of structured ncRNAs (Figure 5C). By contrast, changes in protein-coding gene expression were more heterogeneous, and GO enrichment returned diverse processes such as chromatin organization, signaling and cellular motility rather than a single dominant pathway (Figure 5D, E). With the present data we cannot determine whether these changes are direct or indirect.

**Figure 5.**
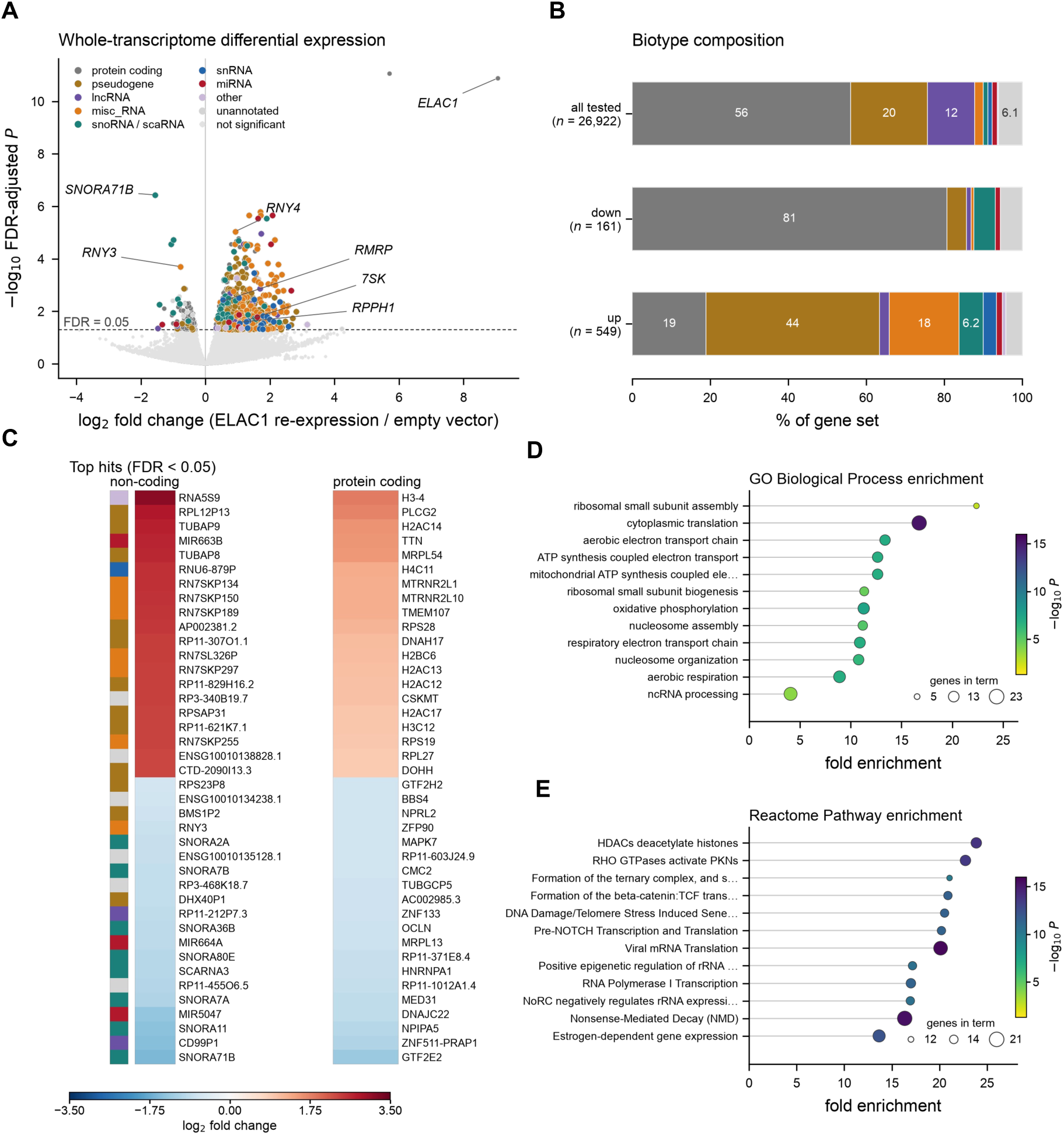
Re-expression of ELAC1 preferentially restores structured non-coding RNAs but not the wild-type transcriptome. Whole-transcriptome (>200 nt, ribodepleted) libraries comparing ELAC1 re-expression with empty-vector knockout cells; n = 2 biological replicates per condition, hits at FDR < 0.05. (A) Volcano of −log10 adjusted P against log2 fold change, colored by GENCODE biotype class; grey, not significant; dashed line, FDR = 0.05. Selected RNA polymerase III-transcribed loci and ELAC1 are labeled. (B) Biotype composition of the tested, down-and up-regulated gene sets, each stacked to 100% in the class order of (A); class enrichment relative to all tested genes was assessed by one-sided hypergeometric test. (C) The 20 largest-magnitude hits in each direction, non-coding (left) and protein-coding (right), ordered by log2 fold change; sidebar, biotype class as in (A). Three off-scale loci (ELAC1, RP11-729L2.2, AC006077.3) are omitted so that the range resolves the remaining hits. (D, E) GO Biological Process (D) and Reactome Pathway (E) enrichment for the differentially expressed protein-coding genes: x-axis, fold enrichment; marker area, number of genes in the term; color, −log10 FDR.

### The 50–100 nt structured non-coding RNA pool responds to ELAC1 perturbation

To assess the contribution of ELAC1 to small RNA homeostasis, we performed small RNA sequencing across three conditions: i) knockout versus wild-type, ii) overexpression versus empty vector, and iii) rescue versus empty vector (Supplementary Figure S4). The 50–100 nt size fraction, capturing full-length and precursor tRNAs, Y RNAs, vtRNAs, and C/D box snoRNAs, was mapped to the human genome using GENCODE annotation. Each condition produced a distinct pattern of differential expression, indicating that ELAC1 levels are associated with ncRNA composition. In the rescue condition, ELAC1 is re-expressed at roughly 530-fold over the empty-vector knockout baseline (log2 fold change +9.05, adjusted P = 1.3 × 10⁻¹¹, total RNA-seq), so this comparison tests the effect of supraphysiological ELAC1 rather than restoration of endogenous levels. No vault RNA locus reaches FDR < 0.05 in this library. Most H/ACA box snoRNAs and several scaRNAs (SCARNA3, SCARNA11, SCARNA16) were downregulated and the majority of snRNAs were upregulated (Supplementary Figure S4A, D).

ELAC1 overexpression produced a broadly reciprocal pattern with snoRNAs and SCARNA3 being among the most upregulated transcripts, while RNY4 and additional snoRNAs declined (Supplementary Figure S4B, E). Rescue of ELAC1 expression in knockout cells caused further rearrangements, with broader changes in snoRNA and tRNA abundance (Supplementary Figure S4C, F). No locus in this comparison passes FDR < 0.05, so it is described only at the class level and no individual loci are named. These results indicate that the response differs by RNA class rather than affecting structured ncRNAs uniformly: in knockout cells H/ACA box snoRNAs and scaRNAs fall while snRNAs rise, and overexpression produces a broadly reciprocal snoRNA response.

### ELAC1 modulates tRNA expression in knockout and rescue conditions

Mapping of small RNA sequencing reads to nuclear and mitochondrial tRNA sequences revealed that ELAC1 perturbation affects both the abundance and composition of cellular tRNA populations. In ELAC1 knockout cells, 84 of 490 tRNA genes were differentially expressed (51 up, 33 down; FDR < 0.05). Mitochondrially encoded tRNAs shifted upward relative to cytosolic ones, 9 of the 12 significant mitochondrial tRNAs being upregulated (Mann–Whitney P = 0.027), which points to distinct regulatory effects across the two pools (Figure 6A, B). At the isodecoder level, GAU-Ile (2 of 3 members, mean log2 fold change −1.6) and UCA-Sec (1 of 2, −1.1) were reduced and CUG-Gln was modestly increased (6 of 18 members, mean +0.3). No member of the UAU-Ile group reached FDR < 0.05 (Figure 6C). Most individual tRNAs followed the trends of their isodecoder groups. Some, however, behaved discordantly, which may reflect alternative processing or differential contributions to tRNA-derived fragment pools. Overexpression of ELAC1 in wild-type cells yielded no tRNA gene passing FDR < 0.05 (0 of 490) (Figure 6D, E), while nominal-level shifts in specific isodecoder groups were still observed (Figure 6F). Elevated ELAC1 therefore appears to perturb tRNA populations less than its complete loss and may act as a regulator without being stoichiometrically limiting. Reintroduction of ELAC1 into knockout cells likewise yielded no tRNA gene at FDR < 0.05 (Figure 6G, H). Because this comparison is rescue against empty-vector knockout rather than against wild type, it asks whether re-expression moves the tRNA pool away from the knockout state, and it does not (Figure 6I). These results suggest that ELAC1 contributes to maintaining the composition of the cellular tRNA pool, its loss producing extensive changes in isodecoder representation that re-expression does not measurably reverse at the level of individual genes.

**Figure 6.**
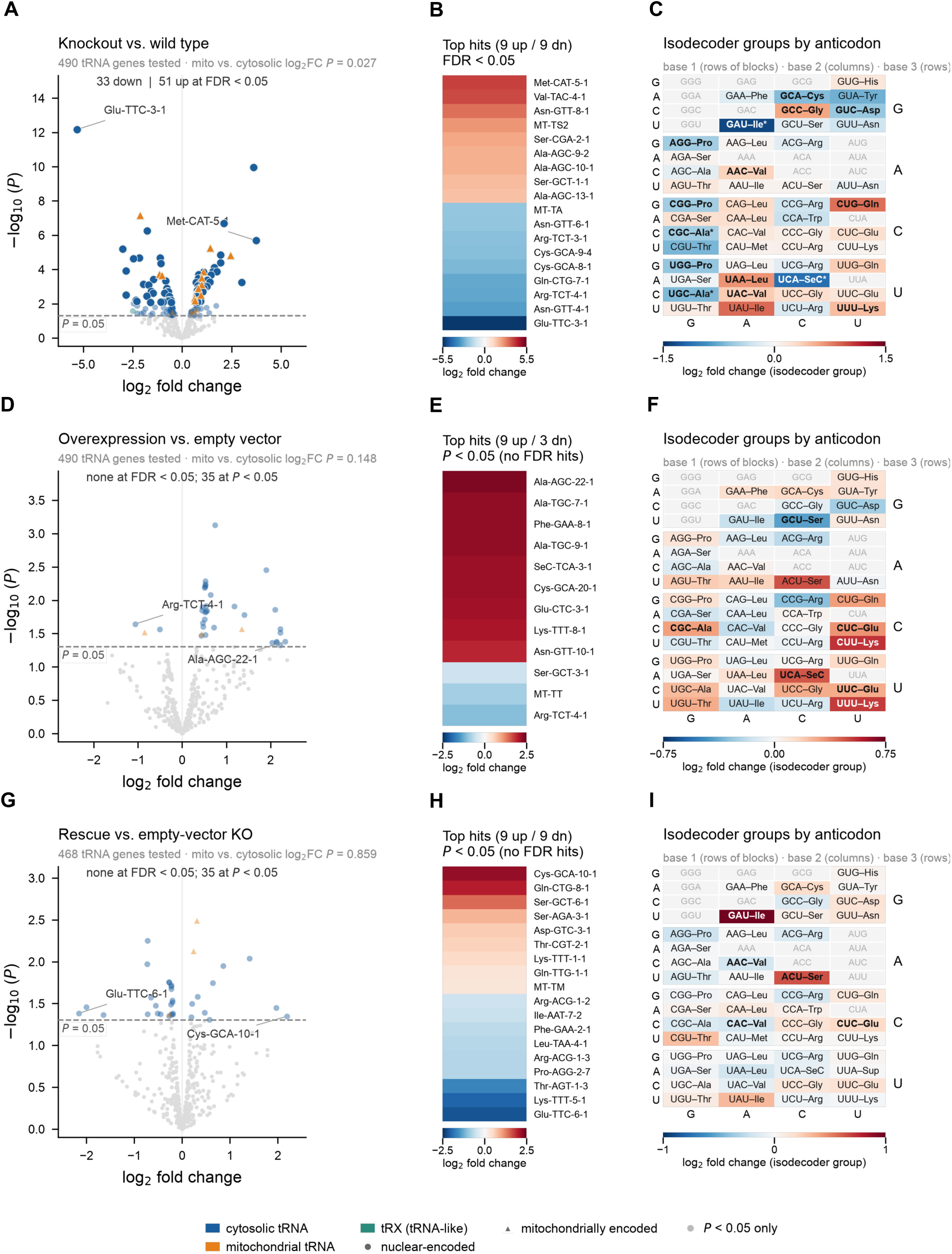
ELAC1 modulates cellular tRNA abundance and isodecoder composition. Rows: ELAC1 knockout versus wild type. (A–C), ELAC1 overexpression versus empty vector (D– F), and ELAC1 re-expression versus empty-vector knockout (G–I). Reads were mapped to nuclear and mitochondrial tRNA genes and tested with limma-voom (n = 2 biological replicates per condition). (A, D, G) Volcano plots of log2 fold change against −log10 nominal P, colored by tRNA gene class; circles, nuclear-encoded; triangles, mitochondrially encoded. Saturated points pass FDR < 0.05, pale points are nominally significant only and grey points are not significant; dashed line, P = 0.05. Each panel header gives the number of genes tested and a two-sided Mann–Whitney test of mitochondrially encoded against cytosolic log2 fold changes. (B, E, H) Log2 fold change of the largest-magnitude hits, up to nine per direction, ordered from most up-to most down-regulated; genes are drawn from the FDR-significant set where one exists and otherwise from the nominally significant set, as stated on each panel. (C, F, I) Isodecoder-group log2 fold changes arranged as a codon table: blocks by anticodon base 1, columns by base 2, rows by base 3; bold, P < 0.05; asterisk, FDR < 0.05. Greyed cells are anticodons absent from the quantification. The three comparisons are independent contrasts, not a matched series.

### ELAC1 level is associated with shifts in tRF class composition and in tRF dependence on precursor tRNA abundance

tRF-1 species are released from the pre-tRNA 3′ trailer, so an enzyme that processes that trailer could in principle shape the tRF pool. Since tRF-1s are generated through RNase Z activity (16), we analyzed tRF abundance and class distribution across all three experimental conditions. The major tRF classes were detected in all conditions, though their relative proportions varied substantially (Figure 7). In ELAC1 knockout cells, 333 of 3,896 tested tRFs were differentially expressed (182 up, 151 down; FDR < 0.05). Upregulated fragments were enriched for 3′-tRFs (46% of hits versus 31% of the tested background). Downregulated fragments were dominated instead by 5′-tRFs (62% versus 19%) and 5′-halves (19% versus 4%), with 3′-tRFs depleted among them (4% versus 31%). Internal tRNA-derived fragments (i-tRFs) were under-represented in both directions (10% and 12% against 37% of the background). ELAC1 loss therefore seems to shift the balance between tRF classes rather than uniformly affecting all fragments (Figure 7A-C). In contrast, overexpression of ELAC1 in wild-type cells produced a much smaller and differently distributed response with 47 fragments found at FDR<0.05, 10 up and 37 down. 3′-tRFs dominated the downregulated fragments (70%), and the ten upregulated fragments comprised three 3′-tRFs, three tRF-1s, two 5′-tRFs and two i-tRFs (Figure 7D-F). Reintroduction of ELAC1 into knockout cells reversed these class-level effects. Across all detected fragments, 5′-tRFs and 5′-halves shifted upward (median log2 fold change +0.40 and +0.21, against −0.43 and −0.76 in the knockout) while 3′-tRFs shifted downward (−0.05, against +0.48). Each class was tested against all remaining fragments (Mann–Whitney *P* = 1.1 × 10⁻¹⁸, 4.6 × 10⁻⁴ and 2.7 × 10⁻⁷). Only eight individual fragments reached FDR<0.05 in this comparison (Figure 7G-I), so the reversal is described at the class level and not as a hit list. Few individual fragments survive multiple-testing correction in the overexpression and rescue comparisons. Both, however, contain substantially more small *P* values, with 283 fragments at nominal *P*<0.05 against 152 expected in the overexpression comparison, and 141 against 95 in the rescue comparison (binomial *P*=4.9 × 10⁻²³ and 3.6 × 10⁻⁶). Both perturbations therefore alter tRF abundance, and across all three conditions the shift is evident at the level of tRF class composition.

**Figure 7.**
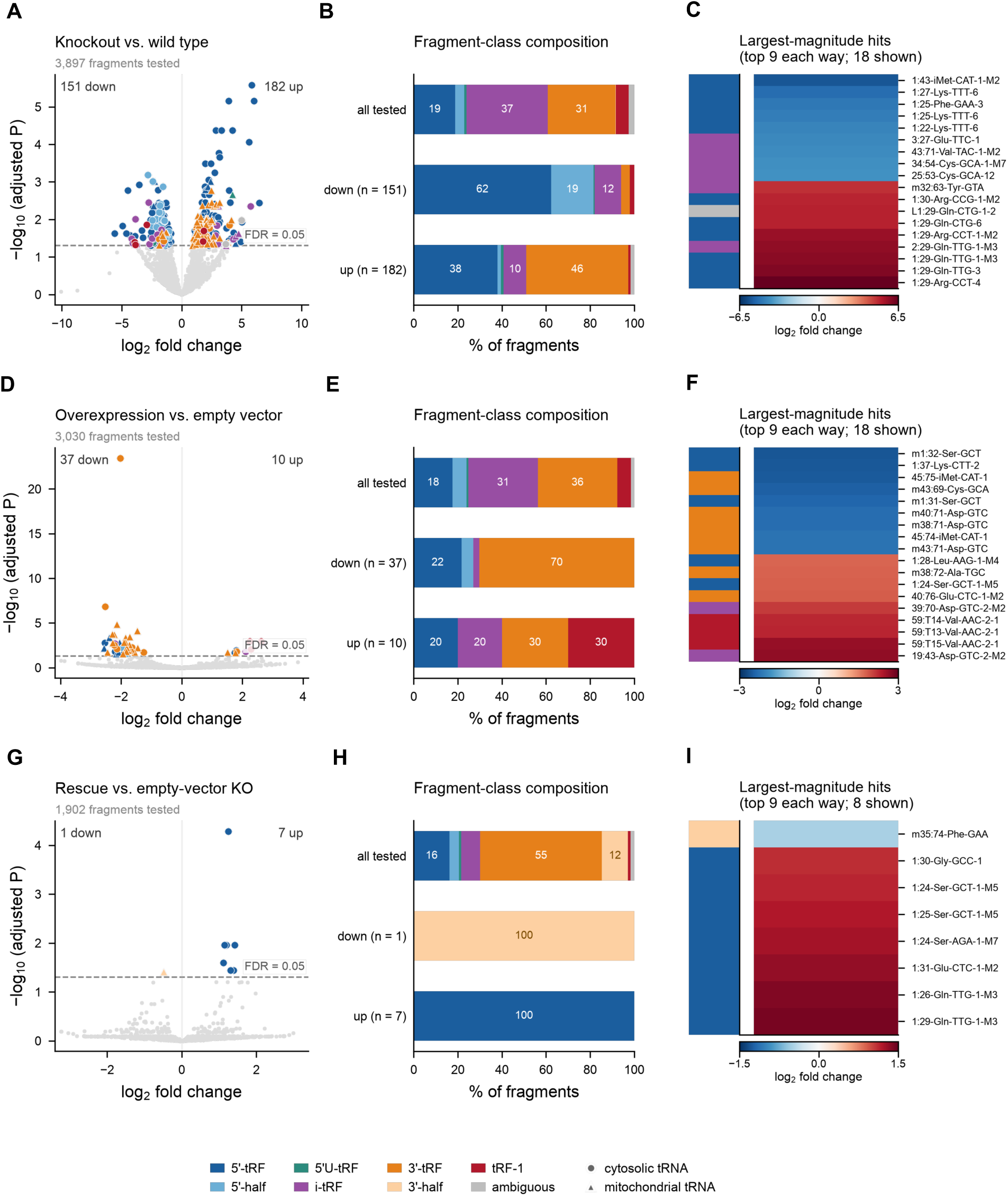
ELAC1 abundance reshapes the composition of tRNA-derived RNA fragments. (A–C) ELAC1 knockout versus wild type; (D–F) ELAC1 overexpression versus empty vector; (G– I) ELAC1 re-expression versus empty-vector knockout; n = 2 biological replicates per condition. (A, D, G) Volcano plots of log2 fold change against −log10 adjusted P; dashed line, FDR = 0.05. Fragments passing FDR < 0.05 are colored by class and non-significant fragments are grey; circles, cytosolic tRNA origin; triangles, mitochondrial. (B, E, H) Class composition of all tested fragments and of the down-and up-regulated sets, as the percentage of each set; the ‘all tested’ row is the detected background against which enrichment is read. (C, F, I) The nine most negative and nine most positive fragments by log2 fold change among those passing FDR < 0.05, with class indicated by the sidebar; only eight fragments reach significance in the re-expression comparison, and all are shown. Fragment names follow MINTmap tDR nomenclature with the ‘tDR-’ prefix omitted; an ‘m’ prefix denotes mitochondrial origin.

### The 15–50 nt small RNA fragment population is redistributed in ELAC1-perturbed cells

To characterize the small RNA populations affected by ELAC1, we analyzed the 15-50 nt fraction from knockout, overexpressing and rescue conditions using SPORTS1.1. This annotates fragment classes beyond tRFs and miRNAs, which include, among others, snoRNA-derived RNAs (snoDRs), snRNA-derived RNAs (snDRs), rRNA-derived RNAs (rDRs), Y RNA-derived RNAs (yDRs) and vtRNA-derived RNAs (vtDRs). Across all conditions, the most broadly affected fragment classes were snDRs and snoDRs. A dedicated miRNA analysis identified 31 differentially expressed miRNAs in knockout cells at FDR<0.05 (12 up, 19 down of 619 tested). Neither the overexpression nor the rescue comparison gave evidence of miRNA changes. No miRNA reached FDR<0.05 in either, and the number passing a nominal *P*< 0.05 cut-off did not exceed chance expectation. For overexpression, 20 miRNAs passed against 31 expected of 619 tested; for rescue, 22 against 17 of 345 (Supplementary Figure S5).

At the level of individual fragment classes (excluding tRNA-and miRNA-derived fragments, which are analyzed in Figures 6, 7 and S5), knockout cells showed the most extensive redistribution. Of the 110 upregulated fragments at FDR < 0.05, rDRs accounted for 49%, snDRs 23%, snoDRs 14% and yDRs 6%; the rDRs derived predominantly from 28S and 18S rRNA and from mitochondrial 12S and 16S rRNA (Figure 8A, D). Among the 76 downregulated fragments, rDRs made up 39% (from 28S, 18S, 5S and 5.8S rRNA), snoDRs 18%, snDRs 17% and piRNA-derived fragments 12%. In overexpressing cells, the response was almost entirely downward (55 of 59 fragments at FDR < 0.05) and dominated by rDRs (85%), with snDRs contributing 11%. Those downregulated snDRs derived from the U1 and U2 snRNA families with RNU6 appearing among them only at nominal significance. The four upregulated fragments were two snDRs, one from the RNU5 family (RNU5E-6P) and one from the RNU1 family, and two 5S rDRs (Figure 8B, E). No fragment reaches FDR<0.05 in the rescue comparison, so no rescue hit list is reported. Figure 8C, F displays that comparison at a nominal *P*< 0.05 cut-off, labeled on the panel, to show the class-level pattern only. vtDRs followed a similar pattern: broadly upregulated in knockout cells and downregulated upon ELAC1 overexpression (Figure 8A). Those derived from VTRNA2-1 were the exception, being undetectable.

**Figure 8.**
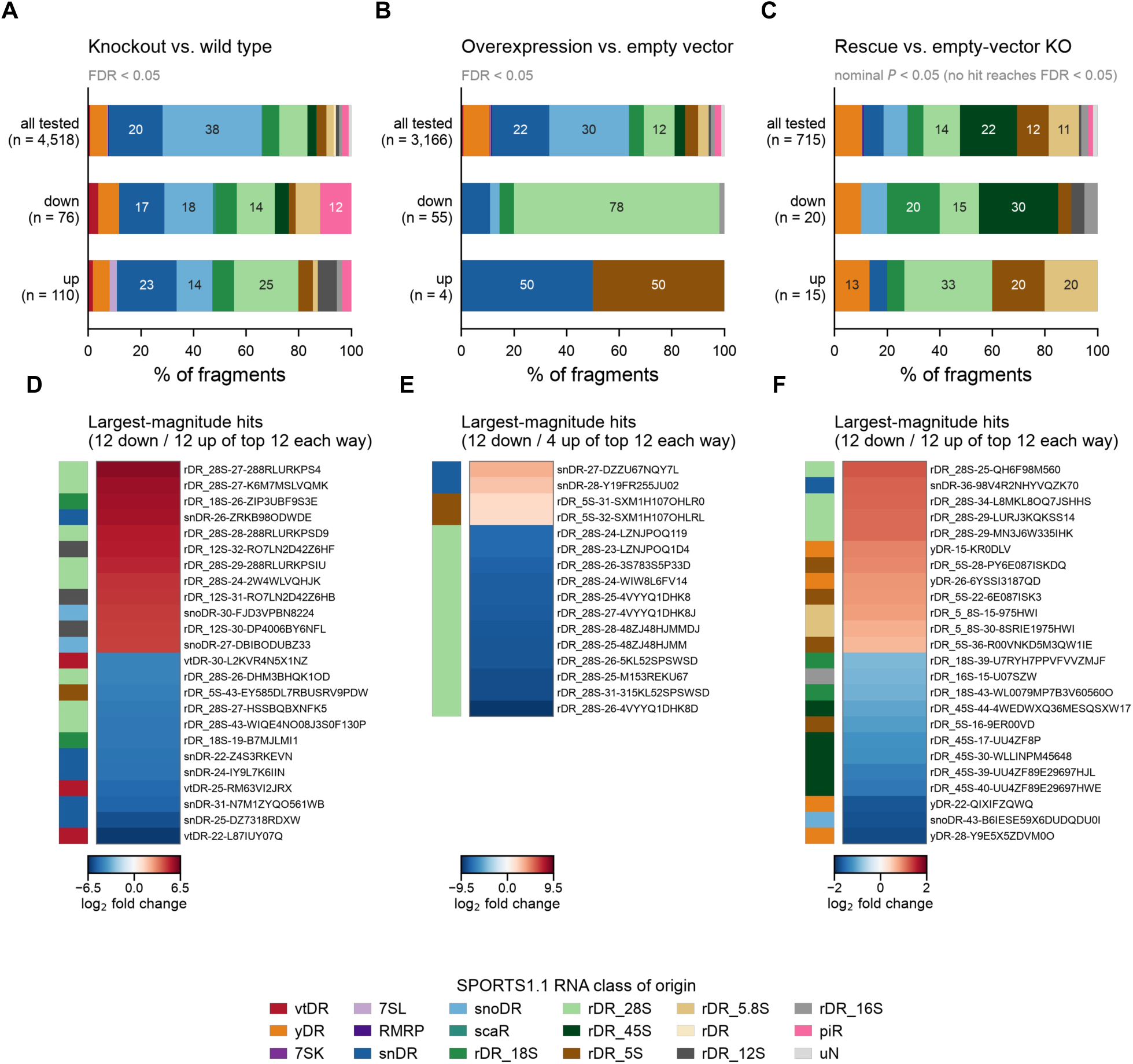
ELAC1 perturbation remodels the composition of small non-coding RNA fragments. Small RNA libraries were annotated with SPORTS1.1 and tested with limma-voom (n = 2 biological replicates per condition). tRNA-derived (tDR) and miRNA fragments are excluded here and are analyzed in Figure 7 and Supplementary Figure S5, respectively; colors denote the SPORTS1.1 RNA class of origin throughout. (A–C) Class composition of the annotated fragment universe (all tested) and of the significantly down-and up-regulated fragment sets, for ELAC1 knockout versus wild type (A), ELAC1 overexpression versus empty vector (B) and ELAC1 re-expression versus empty-vector knockout (C). Stacked bars give the percentage of fragments in each class, with the number behind each bar on the left. Hits are defined at FDR < 0.05 in (A) and (B); no fragment reaches FDR < 0.05 in (C), so that panel uses a nominal P < 0.05 cut-off and its hit sets are descriptive. (D–F) The largest-magnitude hits of each comparison, up to 12 per direction, with the number shown given in each panel header. Rows are ordered by log2 fold change, with the class sidebar as in (A–C).

### Small RNAs with unprocessed 3′ U-rich ends accumulate in ELAC1 knockout cells

The clearest direct footprint of ELAC1 activity was the accumulation of small RNAs bearing a 3′ UUU motif in knockout cells, and their reduction upon ELAC1 overexpression (Figure 9). A similar trend was seen in rescue cells, though reduced read coverage limits interpretation. These species originate from the 3′ ends of RNA polymerase III transcripts that carry a terminal stem with a 3′ extension, adopting stem-like structures reminiscent of pre-tRNA molecules (Figure 9). Their accumulation in the absence of ELAC1 suggests that these transcripts are direct substrates, normally cleaved before further processing by downstream nucleases. In wild-type cells the resulting fragments would then lack the 3′ extension and be annotated as distinct, shorter species.

**Figure 9.**
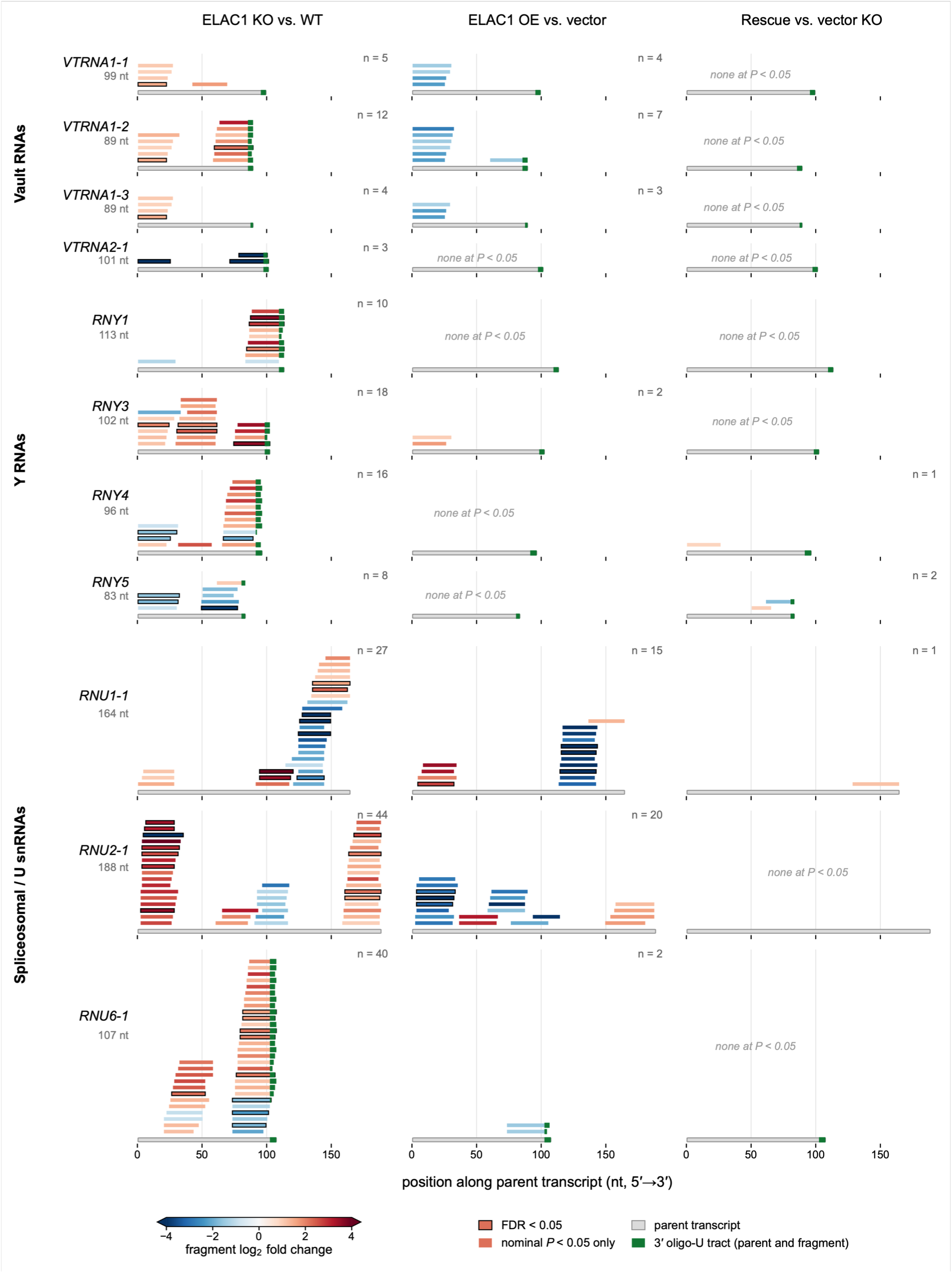
Loss of ELAC1 leads to accumulation of small RNA fragments containing an unprocessed 3′ UUU motif. Rows are parent transcripts (vault RNAs, Y RNAs, spliceosomal / U snRNAs); columns are the three independent comparisons. Each bar is one SPORTS1.1 fragment, matched to the parent by exact sequence match and drawn at its position along it; stacking only prevents overlap, so vertical position carries no meaning. Grey body: the parent transcript, on a nucleotide scale (5′→3′) shared by all cells, with its length beside the row label. Fill: fragment log2 change on one scale centered at zero, clipped at ±4 (color-bar arrowheads). Black outlines mark FDR < 0.05; bars without outlines pass the nominal P < 0.05 inclusion threshold. Green: the genomically encoded 3′ oligo-U tract, on the parent and on fragments retaining part of it (RNU1-1 and RNU2-1 are Pol II and have none).

## Discussion

Our findings identify ELAC1 as a cytoplasmic ribonuclease with a substrate repertoire that extends beyond canonical pre-tRNA processing. ELAC1 associates with, and directly cleaves, a defined set of structured non-coding RNAs, with strong enrichment for RNA polymerase III transcripts and in particular the vault RNAs. Loss and re-expression of ELAC1 are accompanied by widespread alterations in ncRNA abundance and small RNA fragment composition. Because the underlying datasets were not generated as a single matched series, we treat these transcriptome-wide effects as supporting cellular context for candidate substrates rather than proof of direct processing.

The biochemical basis for this expanded substrate repertoire is rooted in fundamental differences in how ELAC1 and ELAC2 engage structurally heterogeneous substrates, as revealed by our analysis of pre-tRNA^Gly^_GCC_. Unlike the well-folded pre-tRNA^Arg^, pre-tRNA^Gly^_GCC_ adopts two conformations in the presence of Mg²⁺, and the two enzymes partition between them. ELAC1, with its substantially lower efficiency on canonical substrates but broader structural tolerance, preferentially acts on the more abundant alternative conformation, cleaving at an internal position within the structured 3′ trailer rather than at the canonical site immediately downstream of the discriminator base. This tolerance for non-canonical conformations raises the possibility that ELAC1 acts on tRNAs that fail to fold correctly, although whether that would salvage them or commit them to turnover is not addressed by our data.

Cryo-EM structures of ELAC2 captured before and after 3′-trailer cleavage define how the trailer is fed into the active site, and structures of the mitochondrial RNase Z complex bound to a CCA-containing tRNA explain the CCA anti-determinant effect (40–43). In ELAC2 the first cytidine is contacted by Ser726 of the conserved HFSQRY motif and the second inserts into a pocket in the N-terminal domain, pulling the scissile bond away from the active site. However, this motif is intact in ELAC1, so the weaker anti-determination of ELAC1 cannot be attributed to its loss. The adjacent PxKxRN motif, which contacts the pre-tRNA in the ELAC2 structures (40) and has been suggested to contribute to CCA anti-determination (44), is only partly conserved. ELAC1 retains the proline and arginine of that motif but substitutes the lysine and asparagine (ELAC2 PMKIRN 493–498; ELAC1 PSPTRG 15–20), and Ile492, whose carbonyl contacts the first cytidine, is a tyrosine. One possibility lies in how the two enzymes assemble their substrate-binding surface: ELAC2 forms it within a single 826-residue chain, whereas in the ELAC1 model an equivalent surface is completed across the interface of the homodimer (Supplementary Figure S6A). Because the second cytidine is read by the N-terminal domain in ELAC2, an equivalent pocket in ELAC1 would have to be contributed by the partner protomer, a prediction that follows from its dimeric architecture. We propose that ELAC1 recognition is not confined to the canonical pre-tRNA fold, extending to structured RNA substrates that present an accessible terminal stem followed by a 3′ single-stranded extension. This organization is found in many RNA polymerase III transcripts, such as vtRNAs and Y RNAs, consistent with our biochemical and CLIP-seq data identifying these as prominent ELAC1 substrates. Consistent with this weaker anti-determination, cytoplasmic ELAC1 can slowly cleave the CCA end from mature tRNAs, potentially destabilizing them (Supplementary Figure S6B).

Loss of ELAC1 results in widespread transcriptome reprogramming, with the most prominent effects on cell adhesion, migration and extracellular matrix remodeling, and weaker enrichment of antiviral response terms. These observations are consistent with prior reports linking ELAC1 expression to interferon responses and viral infection (45, 46), and are compatible with a model in which ELAC1 influences these processes indirectly through its effects on ncRNA processing and stability. Re-expression of ELAC1 in knockout cells does not restore a wild-type transcriptome but instead triggers pronounced upregulation of multiple ncRNA classes. The construct drives ELAC1 far above endogenous levels, so this reflects the properties of the construct as much as the stability of the knockout state. These observations derive from a single knockout clone; validation across additional clones and orthogonal depletion strategies will be needed to separate direct consequences of ELAC1 loss from clonal adaptation.

Our small RNA sequencing analyses reveal that ELAC1 perturbation shapes tRF class composition in each condition tested; because the three datasets were not generated as a single matched dose series, we describe this relationship as condition-dependent rather than strictly graded. Consistent with prior studies (3, 16) we find no evidence that ELAC1 is required for the generation of tRF-1 species such as tRF-1001, reinforcing the view that this activity is mainly attributable to ELAC2.

vtRNAs are the most abundantly represented non-tRNA class in the CLIP-seq data, and the substrate class we characterize most completely. vtRNAs are structural components of the vault ribonucleoprotein complex and have been implicated in multidrug resistance, autophagy regulation, and innate immune signaling (47–50), yet their processing and the regulation of their 3′ ends remain poorly understood. Our biochemical data demonstrate that ELAC1 cleaves vtRNA 3′ ends precisely, immediately after U86 in vtRNA1-2 and at the equivalent position in each of the other three paralogs. That residue is the first of the single-stranded 3′ extension, reminiscent of a discriminator base. Cleavage removes the oligo(U)-containing tail, which could influence RNA stability, localization and interactions with RNA-binding proteins including La-related proteins (Supplementary Figure S6B).

The functional consequences of vtRNA 3′ end processing by ELAC1 are likely to be significant. Recent work has shown that vtRNAs promote RNA virus infection by facilitating cytoplasmic relocalization of hnRNP C and ELAVL1/HuR (51), which enhance viral RNA stability and translation. ELAC1-mediated removal of the 3′ oligo(U) extension could therefore remodel the pool of full-length, pro-viral vtRNAs, positioning ELAC1 upstream of established vtRNA effector pathways. Compensatory expression between the VTRNA1 genes and VTRNA2-1 has also been reported (52), so vtRNA abundance in any one clone reflects locus-level regulation as well as processing. It is also important to distinguish ELAC1-mediated endonucleolytic cleavage from the established TUT4/7–DIS3L2 pathway, which uridylates and degrades cytoplasmic vtRNAs and Y RNAs through a mechanistically distinct route (52, 53).

A similar framework may apply to Y RNAs, which form conserved ribonucleoprotein complexes with Ro60 and La and have been identified as endogenous RIG-I ligands during RNA virus infection (18, 54, 55). We propose that ELAC1-mediated remodeling of the Y RNA 3′ end antagonizes La-mediated protection and promotes transitions toward alternative RNA states, indirectly influencing Y RNA stability, localization, and fragmentation. The persistent downregulation of 3′-end-derived Y RNA fragments across knockout and rescue conditions is consistent with this model. These are distinct from the oligo(U)-bearing 3′ fragments that accumulate in knockout cells, which retain the unprocessed extension, whereas the fragments that decline are those whose ends would have been generated by ELAC1 cleavage. Together this points to a specific role for ELAC1 in Y RNA 3′ end processing and in the biogenesis of Y RNA-derived small RNAs. Y RNA 3′-end fragments are also produced by a structure-dependent cleavage that requires intact RO60 and is only partly attributable to RNase L in human cells, so ELAC1-dependent cleavage and this pathway would need to be separated by end mapping in ELAC1-deficient cells (56).

Analysis of the 15-50 nt fraction reveals that the majority of small RNA fragment changes observed upon ELAC1 perturbation are most parsimoniously explained as indirect downstream consequences of altered parent transcript abundance rather than direct ELAC1 processing. The clearest direct footprint of ELAC1 activity is the accumulation of small RNAs bearing a 3′ UUU motif in knockout cells, derived from the 3′ ends of structured RNA polymerase III transcripts with a terminal stem and 3′ extension. These fragments likely represent substrates that ELAC1 normally cleaves before further processing by downstream nucleases. In wild-type cells the resulting fragments lack the 3′ extension and are annotated as distinct, shorter species; in knockout cells the same nucleases act on unprocessed precursors, producing sequence-shifted fragments with altered annotation and potentially altered function. This primary processor model positions ELAC1 as a gatekeeper that determines the precise sequence boundaries of a subset of cytoplasmic small RNAs, with consequences for their downstream regulatory activity.

Collectively, our findings identify ELAC1 as a ribonuclease that directly cleaves vtRNAs and additional structured RNA polymerase III transcripts in vitro, and that is associated with, and may contribute to, broader remodeling of the cytoplasmic non-coding RNA landscape. Its ability to process vtRNAs, Y RNAs and additional ncRNAs, its association with changes in small RNA populations, and its interactions with components of key ribonucleoprotein complexes place ELAC1 within cytoplasmic RNA processing and post-transcriptional regulation. Its connection to innate immunity in these data is indirect. The antiviral terms that do appear rank well below the cell adhesion and extracellular matrix terms, and are more consistent with ELAC1 acting downstream of or parallel to immune signaling than as an upstream regulator. Whether ELAC1 also shares the protein deAMPylase activity recently reported for bacterial and long-form RNase Z enzymes (57) is untested. These findings expand the functional repertoire of RNase Z enzymes and highlight how cytoplasmic RNA processing activities can be repurposed during evolution to regulate diverse aspects of gene expression.

## ACKNOWLEDGEMENTS

Athanasios-Nasir Shaukat was supported by a scholarship for postgraduate studies from the Bodossaki Foundation. Alexandros Maniatis is a recipient of an “Andreas Mentzelopoulos” Scholarship for PhD studies. Both are gratefully acknowledged.

## DATA AVAILABILITY

RNA-seq and small-RNA-seq data and final analysis files generated in this study have been deposited in the Gene Expression Omnibus (GEO) under accession GSE344921, covering both the whole-transcriptome and small-RNA datasets. CLIP-seq data have been deposited in GEO and the accession number will be released on publication.

