## Supplementary Figures for "An ancestral tRNA maturation enzyme repurposed for cytoplasmic non-coding RNA processing"

**Supplementary Figure S1. ELAC2 processes canonical pre-tRNAs with substantially higher catalytic efficiency than ELAC1.**

(A) Time-course analysis and kinetic parameters for pre-tRNA$^{Arg}$ (*D. discoideum*) processing by ELAC1 and ELAC2. (B) Michaelis–Menten fits for the kinetic analyses shown in (A). (C) Catalytic efficiency ($k_{cat}/K_M$) of ELAC1 and ELAC2 on pre-tRNA$^{Arg}$; $1.36 \times 10^{-3}$ versus $537.5 \times 10^{-3}$ nM$^{-1}$ min$^{-1}$, a 395-fold difference. (D) Secondary structure of pre-tRNA$^{Arg}$ indicating the cleavage site. (E) Cleavage assays of pre-tRNA$^{Gly}$ acceptor-stem length variants (−3 to +3 bp) by ELAC1 and ELAC2. (F) Native PAGE analysis of $Mg^{2+}$-dependent folding of pre-tRNA$^{Gly}_{GCC}$, vtRNA1-2 and pre-tRNA$^{Arg}$.

**Supplementary Figure S2. ELAC1 cleaves additional structured non-coding RNAs in vitro.**

(A) CLIP-seq read coverage of ELAC1-WT and ELAC1-H64A across representative structured non-coding RNA loci: per-base sense-strand coverage in counts per million at a cytosolic tRNA gene, RNU6-1, RNY4, 7SK, RPPH1 and RMRP, with window coordinates (hg19) beneath each track pair. Mean coverage across the plotted window, WT versus H64A: 10,753 versus 4,337 CPM (tRNA-Gly-GCC-1-1, 2.5-fold), 1,976 versus 1,232 (RNU6-1, 1.6-fold), 6,357 versus 3,179 (RNY4, 2.0-fold), 544 versus 401 (7SK, 1.4-fold), 115 versus 154 (RPPH1, 0.75-fold, the one locus higher in the mutant) and 253 versus 134 (RMRP, 1.9-fold). (B,C) In vitro cleavage assays using unlabeled (B) or 5′-32P-labeled (C) RNA substrates.

**Supplementary Figure S3. ELAC1 knockout cells display a mild cell-cycle phenotype.**

(A,B) Cell-cycle analysis of wild-type (A) and ELAC1 knockout (B) cells by propidium iodide staining and flow cytometry. (C) Quantification of cells in G0/G1, S and G2/M phases. (D) Per-gene abundance in ELAC1 knockout versus wild-type cells, as the mean reads per million over the two biological replicates on log axes: 35,435 genes are detected in both conditions (Spearman $\rho = 0.950$ on log10 RPM), and the 4,044 genes detected in wild type only and 3,149 detected in knockout only are drawn in the labeled floor strips instead of being dropped by the log transform; the diagonal marks equality and the flanking lines a four-fold change. (E) Volcano plot of the whole-transcriptome comparison at a stringent cut-off (FDR < 0.001 and |log2 fold change| > 2): of 21,552 tested transcripts, 521 are increased and 1,347 decreased, points are colored by GENCODE biotype and grey points fall below the cut-off, which at this depth still includes transcripts passing FDR < 0.05. Selected loci, including ELAC1, are labeled.

**Supplementary Figure S4. ELAC1 abundance influences the expression of structured RNA polymerase III transcripts.**

(A–C) Volcano plots of the genome-mapped 50–100 nt small RNA libraries, comparing ELAC1 knockout with wild-type cells (A), ELAC1 overexpression with empty vector (B), and ELAC1 re-expression with empty-vector knockout (C). Each point is one annotated locus (limma-voom, n = 2 biological replicates per condition); x, log2 fold change; y, −log10 nominal P. The dashed line marks nominal P = 0.05, points are colored by transcript class (key at bottom), and features passing FDR < 0.05 (Benjamini–Hochberg) are ringed in black. The line above each panel gives the number of features tested, the number passing FDR < 0.05, and a two-sided Mann–Whitney test of the log2 fold-change distribution of RNA polymerase III loci (canonical loci plus Pol III-derived pseudogene copies) against all other tested features. Only the knockout comparison contains FDR-significant features (5 of 696), and Pol III loci show no shift against the remaining features in any comparison (P = 0.21, 0.65 and 0.27). Axes are shared across (A–C). (D–F) The 30 features with the lowest nominal P in the corresponding comparison, ordered from most up- (top) to most down-regulated (bottom); the left-hand bar gives the transcript class and cell color the log2 fold change on a per-panel symmetric scale centered at zero. Feature names in bold followed by an asterisk pass FDR < 0.05. Where two or more annotated loci carry identical count vectors, a single representative row is shown.

**Supplementary Figure S5. ELAC1 perturbation alters miRNA expression.**

(A–C) Volcano plots of mature miRNA differential expression: ELAC1 knockout versus wild type (A), ELAC1 overexpression versus empty vector (B) and ELAC1 re-expression versus empty-vector knockout (C). miRNAs at nominal P < 0.05 (dashed line) are colored by hairpin arm (3p, 5p, or no arm suffix in miRBase) and miRNAs below that threshold are grey; features passing FDR < 0.05 are ringed in black and the four lowest-P miRNAs per panel are named. Panels share both axes, and the line above each panel gives the number of miRNAs tested and the number passing FDR < 0.05. Only the knockout comparison contains FDR-significant miRNAs (31 of 619); the overexpression (0 of 619, smallest adjusted P = 0.88) and re-expression (0 of 345, smallest adjusted P = 0.50) comparisons contain none, so panels (B) and (C) show no significant change after multiple-testing correction. (D–F) Log2 fold change of the 30 lowest-P miRNAs in the corresponding comparison, ordered from most up- to most down-regulated; the arm is read from the -3p/-5p suffix of each row label, color limits are per panel, and asterisks with bold labels mark FDR < 0.05. limma-voom, n = 2 per condition

**Supplementary Figure S6. ELAC1 and ELAC2 share the RNase Z catalytic core but differ in domain organization, quaternary structure and substrate range.**

(A) AlphaFold 3 model (58) of the human ELAC1 homodimer bound to an 83-nt pre-tRNA$^{Arg}$(ACG) (left; protomer A, pale green; protomer B, deep green) and human ELAC2 bound to mitochondrial pre-tRNA$^{Tyr}$ (right; PDB 8Z1G; N-terminal domain 52–470, pale red; C-terminal domain 471–793, deep red), both shown in the same orientation after superposition on the metal-binding residues; flexible arm, gold; substrate tRNA, grey with the 3′ trailer in orange; catalytic $Zn^{2+}$, grey spheres. Superposition places ELAC1 protomer A on the ELAC2 N-terminal domain and protomer B on the C-terminal domain (261 of 363 Cα within 4 Å in each case), with neither protomer overlaying the other domain, indicating that ELAC1 reconstitutes across its dimer interface the two-lobed substrate channel that ELAC2 builds within a single polypeptide. (B) Working model for the division of labor between the two paralogs. In the nucleus, ELAC2 removes the 3′ trailer from pre-tRNAs and matures tRNA-like RNAs like mascRNA (59). In the cytoplasm, ELAC1 can potentially remove the CCA from mature tRNAs, acts on tRNAs released from stalled ribosomes, and removes the 3′ UUU tail of free vault RNAs. This removal may block recognition by La-related proteins and other 3′ end-binding factors and thereby favor Dicer processing into vault RNA-derived small RNAs. In the mitochondrion, tRNAs punctuating the polycistronic transcript are excised by mtRNase P and ELAC2. Dotted lines indicate hypothetical paths rather than established routes. Vault particles are not restricted to the nuclear envelope; only that location is drawn here, for simplicity.

### Supplementary Figure S1

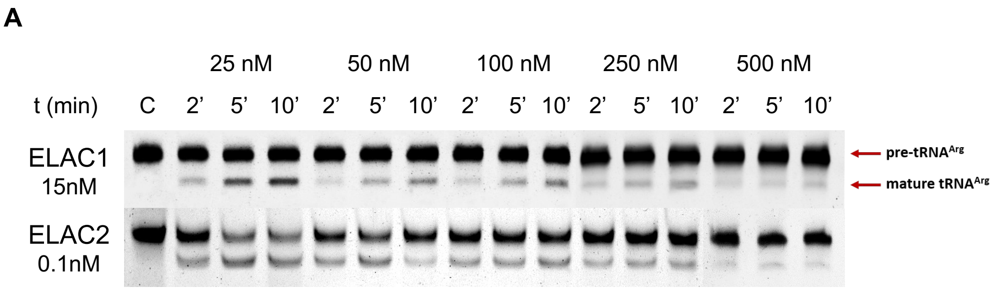


| Best-fit values | ELAC1 | ELAC2 |
|---|---|---|
| [Enzyme] (nM) | 15 | 0.1 |
| Kcat ($min^{-1}$) | 0.6452 | 86.92 |
| Km (nM) | 471.1 | 161.7 |
| Vmax (nM*$min^{-1}$) | 9.678 | 8.692 |
| Kcat/Km | 1.36*$10^{-3}$ | 537.5*$10^{-3}$ |
| **95% CI (profile likelihood)** | | |
| kcat | 0.5571 to 0.772 | 68.42 to 118 |
| Km | 362.4 to 633.5 | 86.56 to 321.8 |
| Goodness of Fit | | |
| **Degrees of Freedom** | 3 | 3 |
| R squared | 0.9991 | 0.9856 |
| Sum of Squares | 0.0127 | 0.2832 |
| Sy.x | 0.06506 | 0.3072 |

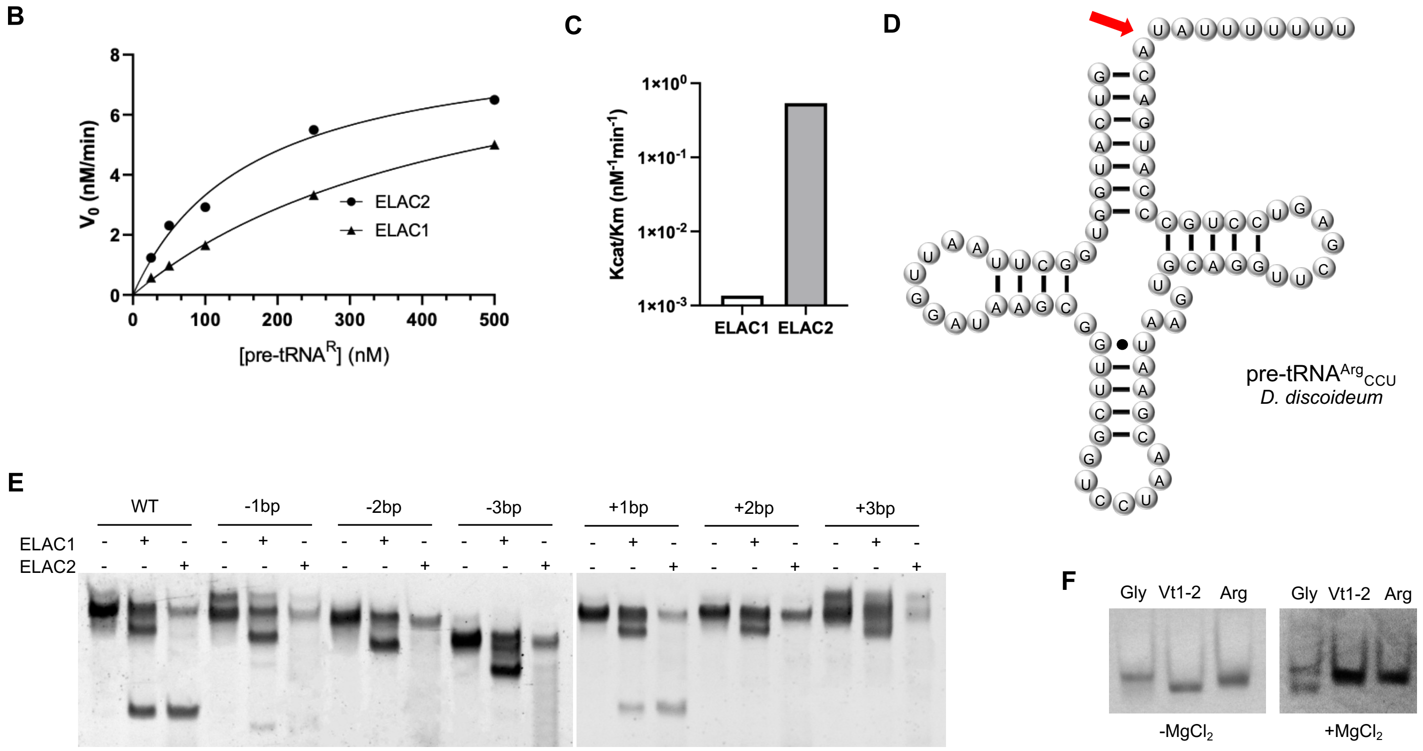

**Supplementary Figure S2**

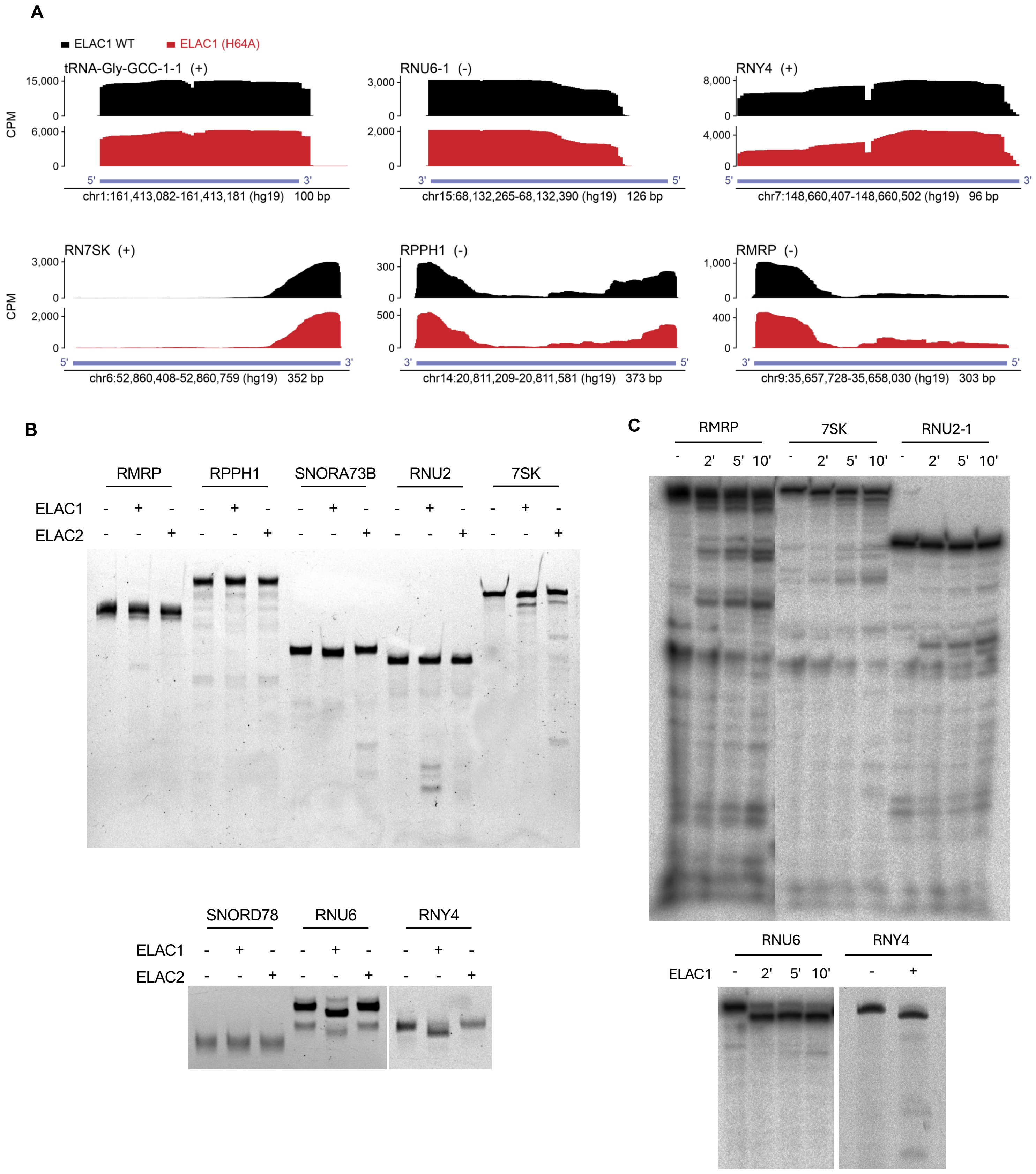

#### Supplementary Figure S3

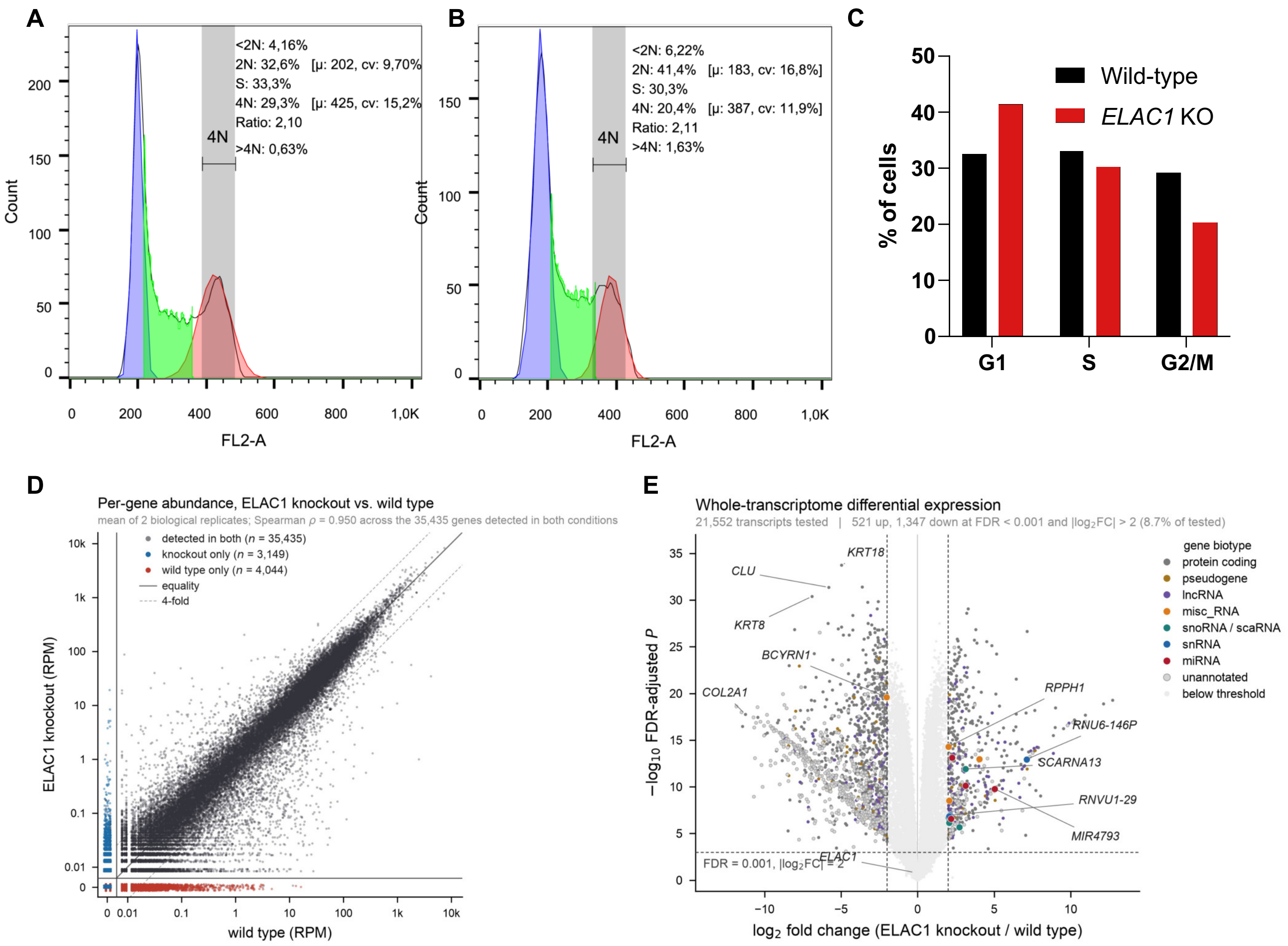

**Supplementary Figure S4**

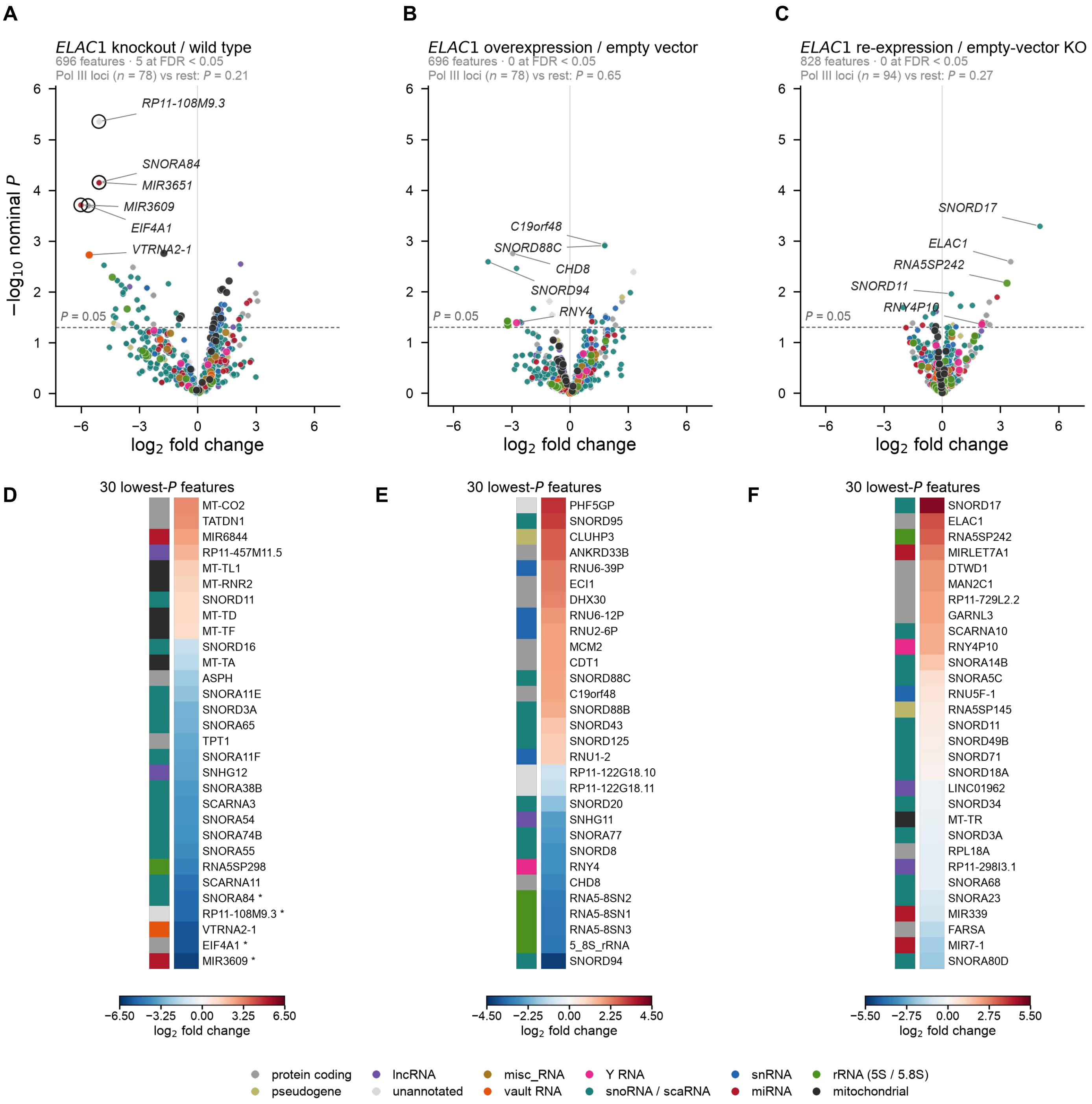

**Supplementary Figure S5**

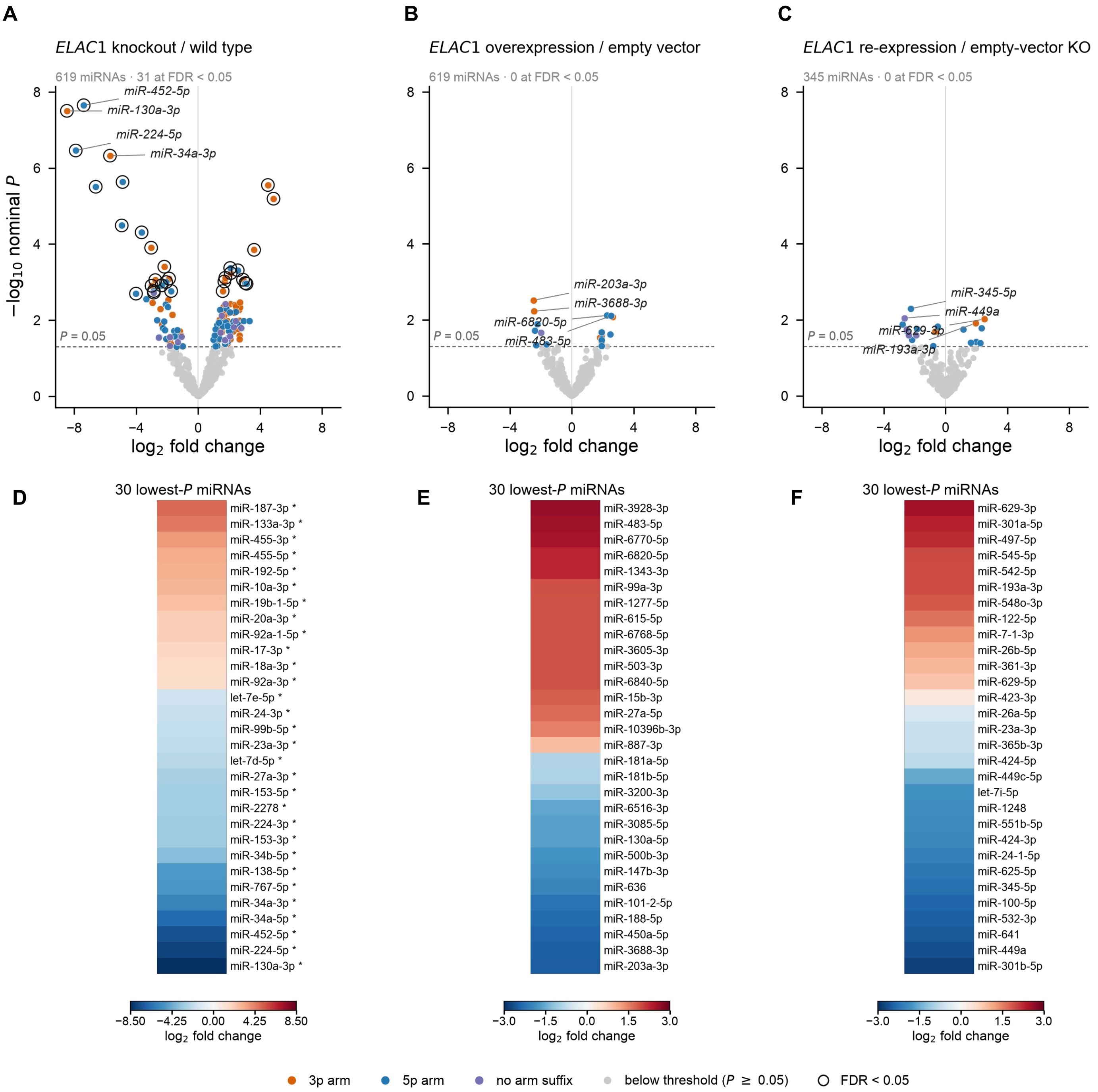

#### Supplementary Figure S6

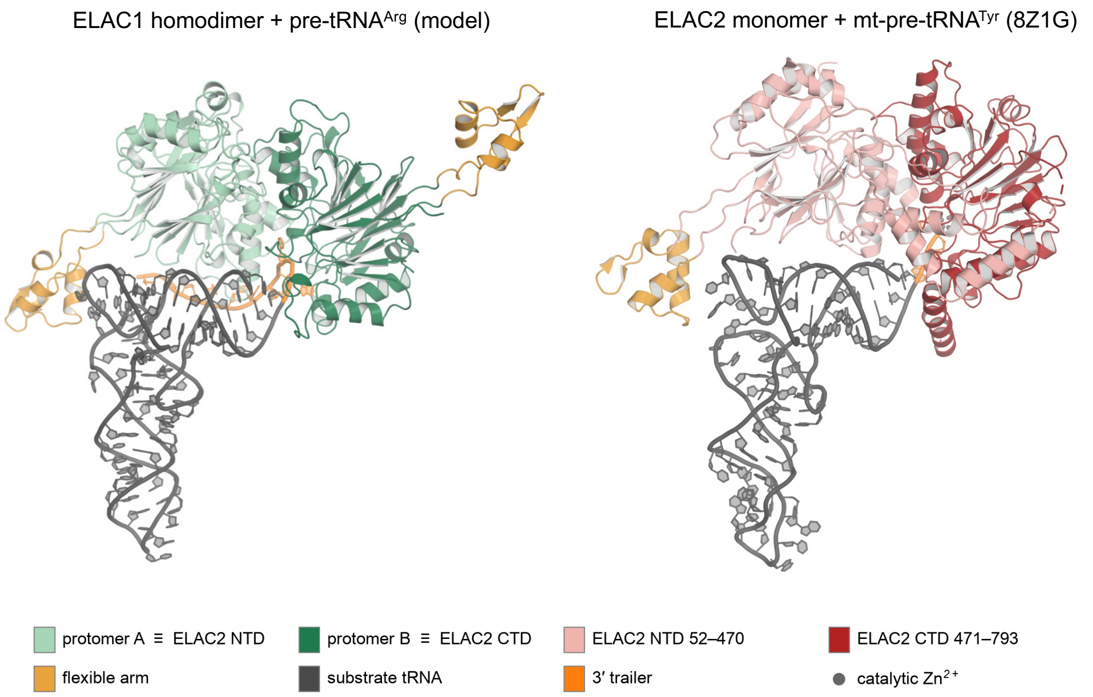


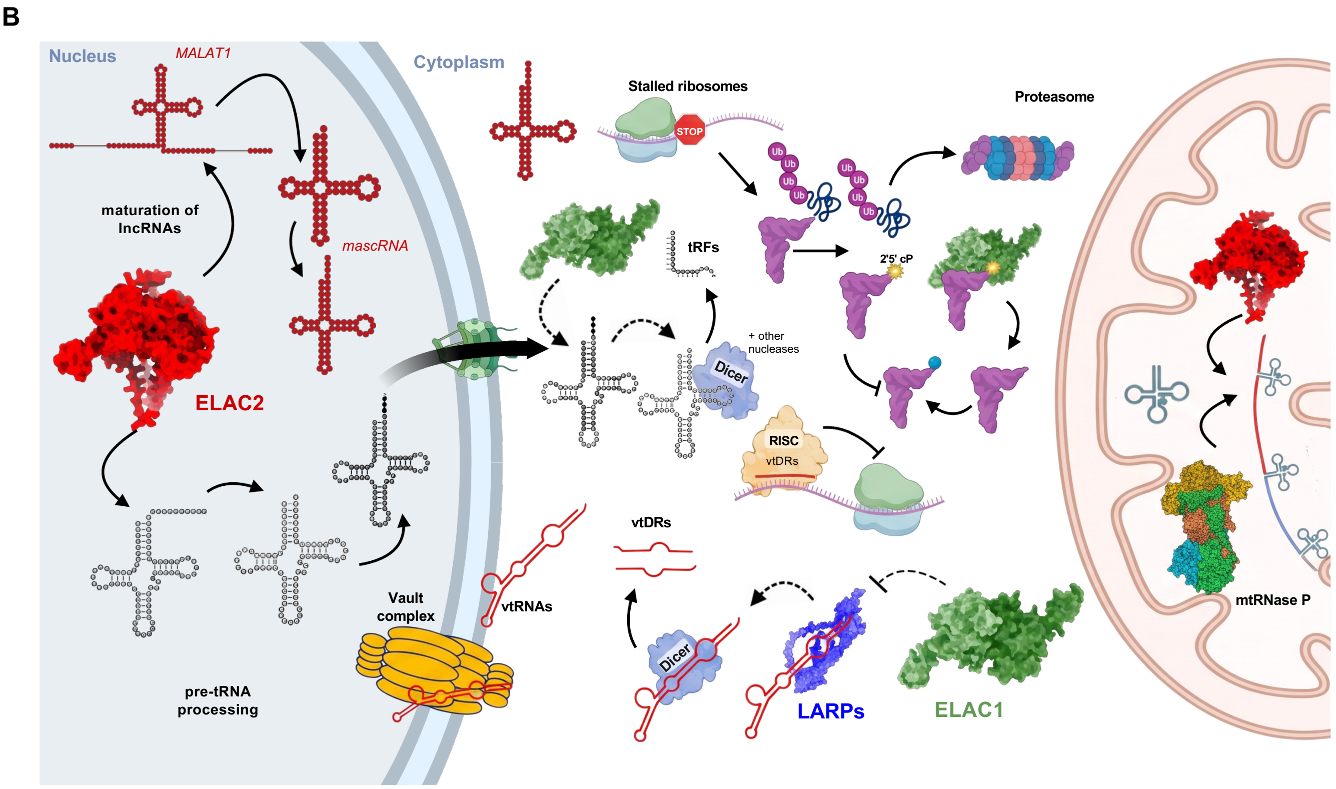
